# Novel temporal and spatial patterns of metastatic colonization from rapid-autopsy tumor biopsies

**DOI:** 10.1101/2021.06.03.446803

**Authors:** Xiaomeng Huang, Yi Qiao, Samuel W. Brady, Rachel E. Factor, Erinn Downs-Kelly, Andrew Farrell, Jasmine A. McQuerry, Gajendra Shrestha, David Jenkins, W. Evan Johnson, Adam L. Cohen, Andrea H. Bild, Gabor T. Marth

## Abstract

**Background:** Metastatic breast cancer is a deadly disease with a low 5-year survival rate. Tracking metastatic spread in living patients is difficult, and thus poorly understood.

**Results:** Via rapid autopsy, we have collected 30 tumor samples over 3 timepoints and across 8 organs from a triple-negative metastatic breast cancer patient. The large number of sites sampled, together with deep whole genome sequencing and advanced computational analysis, allowed us to comprehensively reconstruct the tumor’s evolution at subclonal resolution. The most unique, previously not reported aspect of the tumor’s evolution we observed in this patient was the presence of “subclone incubators”, i.e. already metastatic sites where substantial tumor evolution occurred before colonization of additional sites and organs by subclones that evolved at the incubator site. Overall, we identified four discrete waves of metastatic expansions, each of which resulted in a number of new, genetically similar metastasis sites that also enriched for particular organs (e.g. abdominal vs bone and brain). The lung played a critical role in facilitating metastatic spread in this patient: the lung was the first site of metastatic escape from the primary breast lesion; subclones at this site were the source of all four subsequent metastatic waves; and multiple sites in the lung acted as subclone incubators. Finally, functional annotation revealed that many known driver or metastasis-promoting tumor mutations in this patient were shared by some, but not all metastatic sites, highlighting the need for more comprehensive surveys of a patient’s metastases for effective clinical intervention.

**Conclusions:** Our analysis revealed the presence of substantial tumor evolution at metastatic incubator sites, with potentially important clinical implications. Our study demonstrated that sampling of a large number of metastatic sites affords unprecedented detail for studying metastatic evolution.

## Background

Metastatic breast cancer (MBC) is a deadly disease with a median survival of only 38 months[1]. A previous study estimated that three out of four patients initially diagnosed with stage I-III disease progressed to MBC[2]. Although the genomic and transcriptomic properties of primary tumors have been described extensively[3–5], metastatic tumors, as well as the processes leading to metastasis, are poorly understood because comprehensive biopsying of metastatic sites is difficult or impossible in living patients. Rapid autopsy programs, in contrast, offer pathologists a comprehensive spatial understanding of the extent of the disease, and allow for the collection of fresh tissue samples across all affected organs, within hours of the patient’s death. This approach has been used to study metastatic tumor evolution in breast cancer with TNBC patients being a smaller subset[6–9] and in other cancer types[10–14]. For example, Savas *et al*. studied tumor evolution in 3 estrogen-receptor (ER)-positive, human epidermal growth factor receptor 2 (HER2)-negative breast cancer patients, and 1 triple negative breast cancer patient, using primary tumor and 5-12 matched metastatic samples from the CASCADE program[8]; and Hoadley et al. profiled primary tumors with 4-5 matched metastases genomically and transcriptomically in 2 triple negative breast cancer patients[7]. More recently, De Mattos-Arruda et al. profiled 7-26 samples per patient from autopsies of 10 patients (5 ER+/HER2-, 3 ER+/HER2+, 1 ER-/HER2+, 1 TNBC) with therapy-resistant breast cancer[9]. These studies found significant heterogeneity in both the primary and metastatic tumors, and complex evolutionary patterns during disease progression. However, critical questions remain unanswered, especially in TNBC patients: for example, whether the ability for the cancer to metastasize fully develops in the primary tumor, as suggested by studies[7], or if early metastatic sites can provide niches where the cancer can further develop metastatic potential not present in the primary tumor, but necessary to invade additional organs. To understand metastatic tumor evolution and disease progression at subclonal resolution, we studied the primary tumor at diagnosis and at surgery, as well as 28 metastatic samples across seven organs from a metastatic breast cancer patient with aggressive disease, collected via rapid autopsy following the patient’s death, 30 samples in total. Deep whole genome sequencing allowed us to reconstruct detailed subclone structure and track subclonal expansion across these samples, elucidating the order and timing in which each metastatic site was established, including metastatic colonization events from one organ to another.

## Results

### 1. Clinical presentation showed extremely aggressive metastatic cancer

We studied a 45-year-old woman with ER-negative, PR-negative, HER2-negative metaplastic Grade III invasive ductal carcinoma of the breast (***Fig. 1A***). At the time of diagnosis, she had a clinical T2N0M0 breast cancer, with staging including an ultrasound (US) and MRI showing a 3.1 cm mass. However, the sentinel node biopsy showed none of the three biopsied lymph nodes had cancer. The patient received neoadjuvant therapy of doxorubicin and cyclophosphamide (AC) followed by 8 weeks of weekly paclitaxel. An ultrasound image after the AC showed enlargement of the breast mass. After paclitaxel, the mass remained stable in size on ultrasound but was more painful. The patient underwent mastectomy at week 21. At week 39 after diagnosis, MRIs showed brain metastases, and CT scans showed multifocal metastases in the lung, liver, pancreas, bone, skin, and lymph nodes. This led to subsequent courses of chemotherapy and radiation therapy, all without response, and 56 weeks after diagnosis, the patient succumbed to the disease. Rapid autopsy was performed approximately two hours after death: 28 metastatic tumor samples, 14 surrounding normal tissue samples, as well as 2 normal skin samples were collected (see ***Supplementary Table 1*** for detailed sample descriptions). We were also able to acquire formalin-fixed paraffin-embedded (FFPE) primary breast tumor samples collected at diagnosis (BrP) and at mastectomy (BrM). All tumor samples and 2 normal skin samples were subjected to whole genome sequencing (WGS). All samples except the FFPE samples and the two normal skin samples were also subjected to bulk RNA sequencing. We hypothesized that genomic and transcriptomic analysis of this unprecedented collection of 46 biopsy samples from a single patient would provide the ability to reconstruct the evolutionary course of this aggressive metastatic cancer. According to the wishes of the patient and family, in order to honor the contribution of the patient, and after Institutional Review Boards (IRB) approval, we named this study “The Victoria Clark Study”.

**Figure 1.**
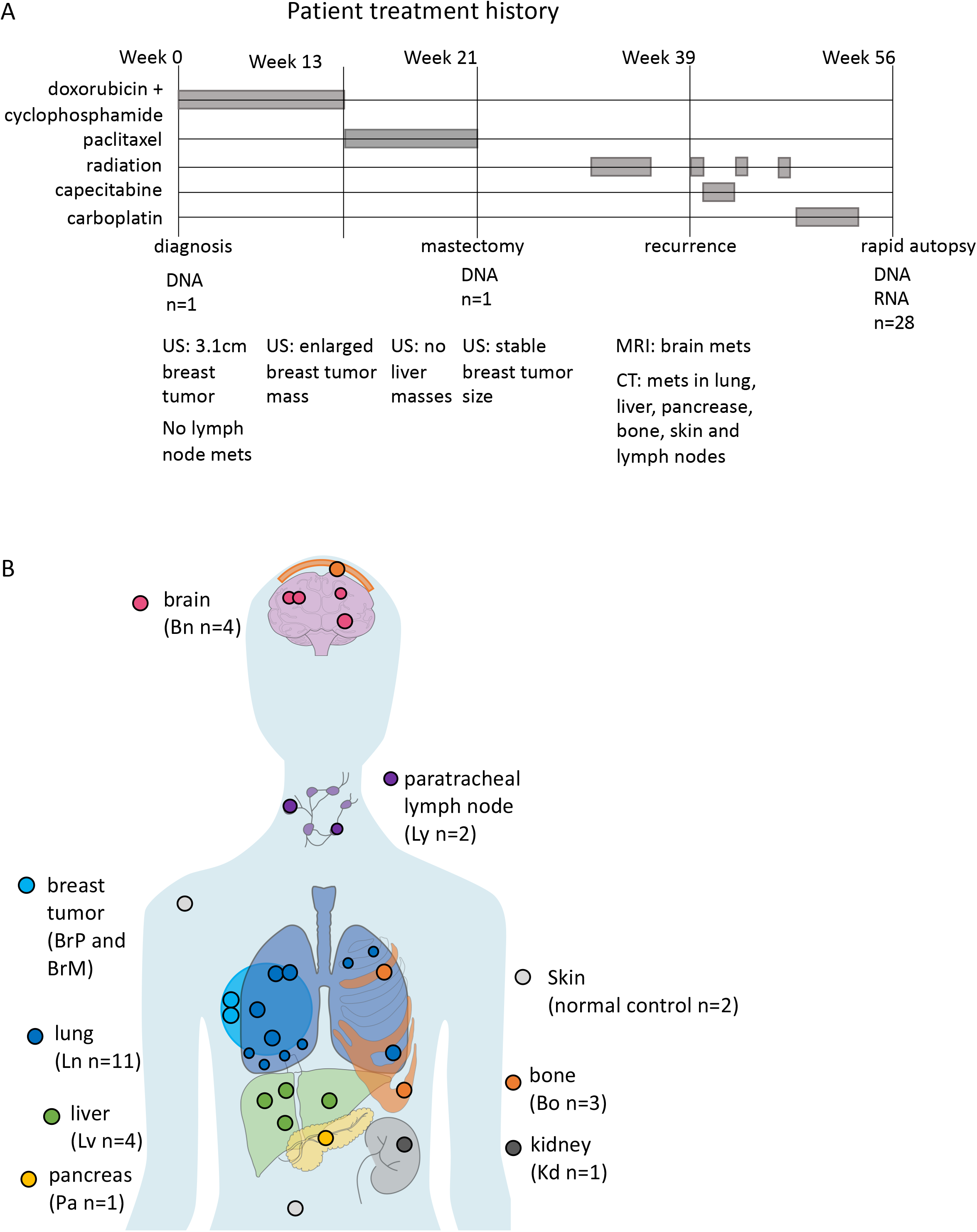
Patient treatment history and sample origins. **(A)** Treatment history over the course of disease progression as well as imaging history including ultrasound (US) imaging, MRI and CT scan. **(B)** Primary breast tumor at diagnosis (BrP) and at mastectomy (BrM) were biopsied. 26 metastatic tumors and 2 normal skin autopsy samples were collected through a rapid-autopsy procedure. Placements of samples are indicative of organ-of-origin, not actual sample locations.

### 2. Genomic characteristics show widespread somatic mutations across all tumor sites

We interrogated the WGS data (∼60X coverage from the rapid autopsy fresh tissue samples, and ∼45X coverage from the FFPE samples, with no discernible quality difference between 60X and 45X samples; see ***Supplementary Fig. S1***) with our state-of-the-art multi-sample tumor data analysis pipeline (see **Methods**), identifying inherited and somatically acquired variants including single nucleotide variants (SNVs), short insertions/deletions (INDELs), copy number variations (CNVs), regions of loss-of-heterozygosity (LOH), and chromosomal translocation events. Tumor purity was estimated by FACETS[15]; 28 tumor samples that had >50% tumor content (see ***Fig. 1B***) were selected for subsequent genomic analysis. The patient had no identifiable germline breast cancer predisposition variants, in concordance with earlier clinical testing for *BRCA1* and *BRCA2* germline mutations. However, all tumor samples, including the primary and mastectomy, showed widespread chromosomal aberrations, including CNVs and large regions of LOH (***Supplementary Fig. S2***). We also found a total of 20,012 somatic mutations across all samples. 5,149 of these were shared by all samples, including a homozygous *TP53* missense (c.517G>C, p.V173L) SNV, a homozygous *PTEN* frameshift (c.676_697delTCCTCCAATTCAGGACCCACAC, p.S226fs), and a homozygous *RB1* deletion; 11,246 additional mutations were shared by at least two samples, and 3,617 variants were sample-specific. The average number of mutations was 10,648 per sample (range 9,975-12,548, see ***Supplementary Table 2***). The numbers of mutations shared among samples vary partly due to the presence of LOH. For example, samples in G1 retained both alleles on chromosome 3 whereas samples in G2, G3, and G4 had lost one of the alleles. All somatic mutations on the lost allele are therefore absent in G2, G3, and G4 samples (***Supplementary Fig. S1, Fig. 2C***). A high fraction, on average 80% per sample, were already present in the primary tumor BrP i.e. these were truncal mutations. 18% were shared mutations with at least one other sample, and the remaining 2% were sample-specific mutations.

**Figure 2.**
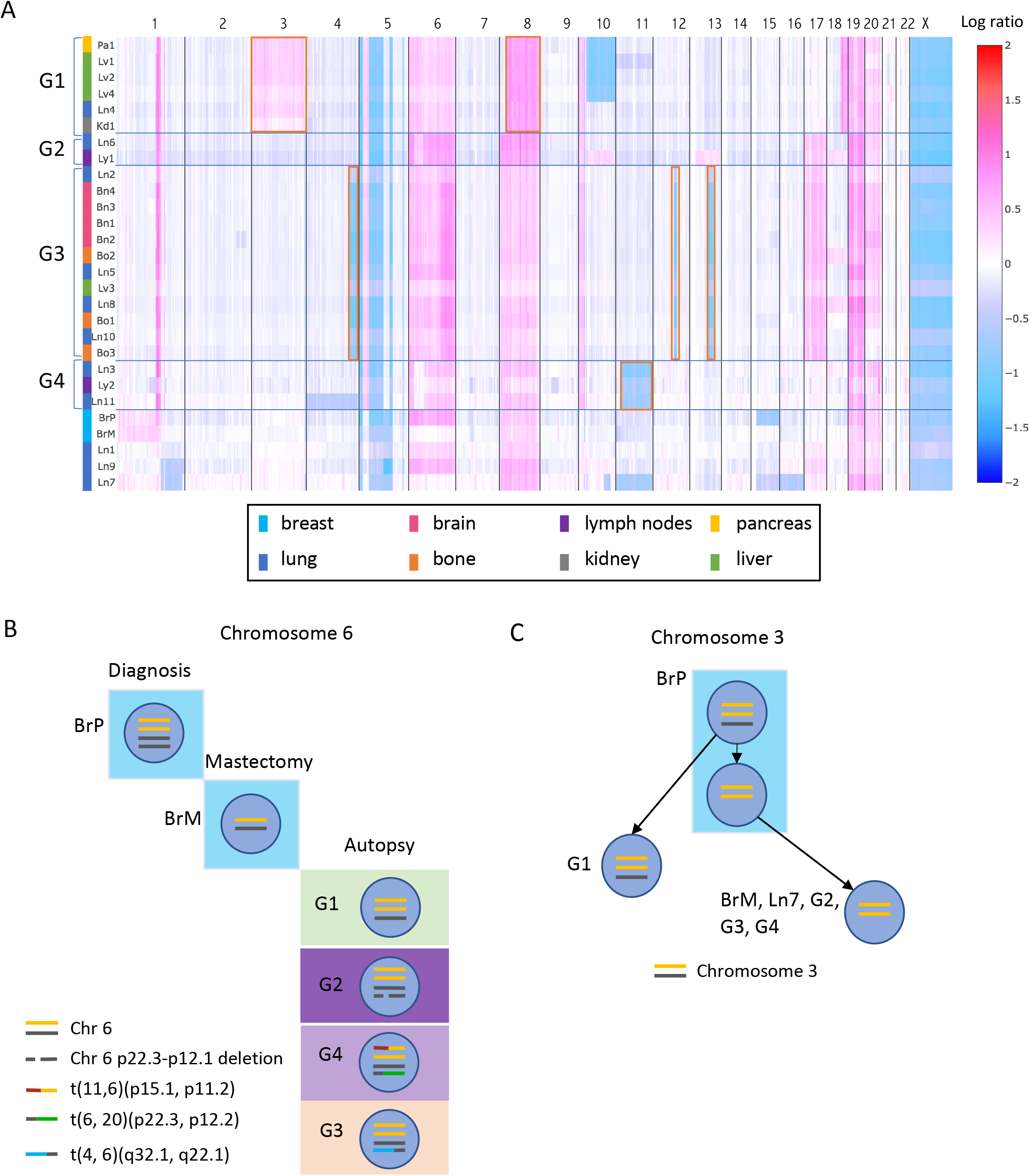
Genomic CNV profiles across samples reveal metastatic waves. **(A)** Heatmap of copy number profile across all samples. Chromosomal coordinates are on the x axis and samples are plotted along the y axis. Colors in the heatmap represent the log2 ratio of tumor copy number to normal copy number. Samples with similar CNV profiles are grouped together as G1, G2, G3, and G4. The shared CNVs in each group are highlighted in red boxes. Colored bars beside the sample names represent the represent the host organs: lung(blue), bone(orange), brain(pink), breast(cyan), liver(green), kidney(grey), pancreas(yellow). **(B)** An example of samples in the same group sharing complicated structural variants on chromosome 6, further confirming CNV based grouping. Yellow and black lines are the two alleles of chromosome 6. A deletion occurred between 6p22.3 and 6p12.1 on one black allele in G2. A translocation occurred between chromosome 11 and one yellow allele of chromosome 6, designated as t(11,6)(p15.1,p11.2)(red and yellow line), and a translocation occurred between one black allele of chromosome 6 and chromosome 20, designated as t(6, 20)(p22.3, p12.2) (black and green line), in G4. A translocation occurred between chromosome 4 and one black allele of chromosome 6, designated as t(4, 6)(q32.1, q22.1) (blue and black line), in G3. **(C)** Copy number changes on chromosome 3 in different groups and inferred evolution. Yellow and black lines are two alleles of chromosome 3. G1, G2, G3, and G4 represent samples in each group. Each blue circle represents a cell population with certain chromosomal features inside.

### 3. Chromosomal changes suggest four distinct waves of metastatic colonization

Based on the similarity of their copy number profiles (***Fig. 2A***), we were able to group 23 of the 28 tumor samples into 4 distinct groups (the shared CNVs in each group are highlighted in red boxes in ***Fig. 2A***). Samples in each group fell into almost perfectly delineated organ groups within the body (G1: abdominal organs, G2: lymph nodes, G3: brain and bones, and G4: lymph nodes). Notably, every group also contains lung sites. Each group signifies a distinct wave of metastatic colonization, as samples in the same group share a common genetic origin. The CNV-based grouping was confirmed by detailed structural variant (SV) analysis (see **Methods**) in which we identified the exact deletion and amplification breakpoints and the specific deleted or amplified alleles (***Fig. 2B***).

We then attempted to reconstruct the time order of these metastatic waves. In contrast to longitudinally sampled cancer genomes with inherent time course, rapid autopsy datasets are collected at a single time point; therefore, the time order in which these sites were established must be inferred from the data. Here we were able to use the observed chromosomal changes to infer partial time ordering: samples in G1 all have one chromosome 3 allele amplified, while still retaining the second allele; whereas samples in groups G2 – G4 have two copies of the first allele but lost the second allele (***Fig. 2C***). This indicates that tumor sites in G1 were seeded by an earlier tumor subclone than sites in the other groups, therefore G1 was likely the first metastatic wave. Furthermore, the breast biopsy collected at mastectomy (sample BrM) consists entirely of the G2-G4 genotype, whereas the primary breast tumor (BrP) is a mixture of cells, containing both the earlier and the later genotypes (***Fig. 2C; Supplementary Fig. S3***). This indicates that a clonal sweep of the G2-G4 lineage occurred in the primary tumor after a G1 precursor escaped. CNV data alone was insufficient to determine the relative time order of the three later waves (G2-G4) and the relationship between the four groups and three lung metastasis samples (Ln1, Ln9, and Ln7) which have distinct copy number profiles and therefore could not be placed in any group (***Fig. 2A***).

### 4. Phylogenetic analysis based on somatic tumor SNVs and short INDELs confirms and refines the four metastatic waves

To further resolve the evolutionary trajectory of the tumor, we constructed a phylogenetic tree among the tumor biopsy samples using somatic SNVs and INDELs (see **Methods**). To avoid any confounding effects of CNV and/or LOH on calculating evolutionary distances between samples (e.g. deletions/LOH events cause samples to lose acquired somatic alleles that reside on the deleted chromosome, resulting in falsely reduced evolutionary distance), we restricted our analysis to somatic mutations on chromosome 14, the only chromosome in our dataset that remained copy number neutral and unaffected by LOH across all tumor samples (***Supplementary Fig. S2***). ***Fig. 3A*** shows the inferred phylogenetic relationship across all samples.

**Figure 3.**
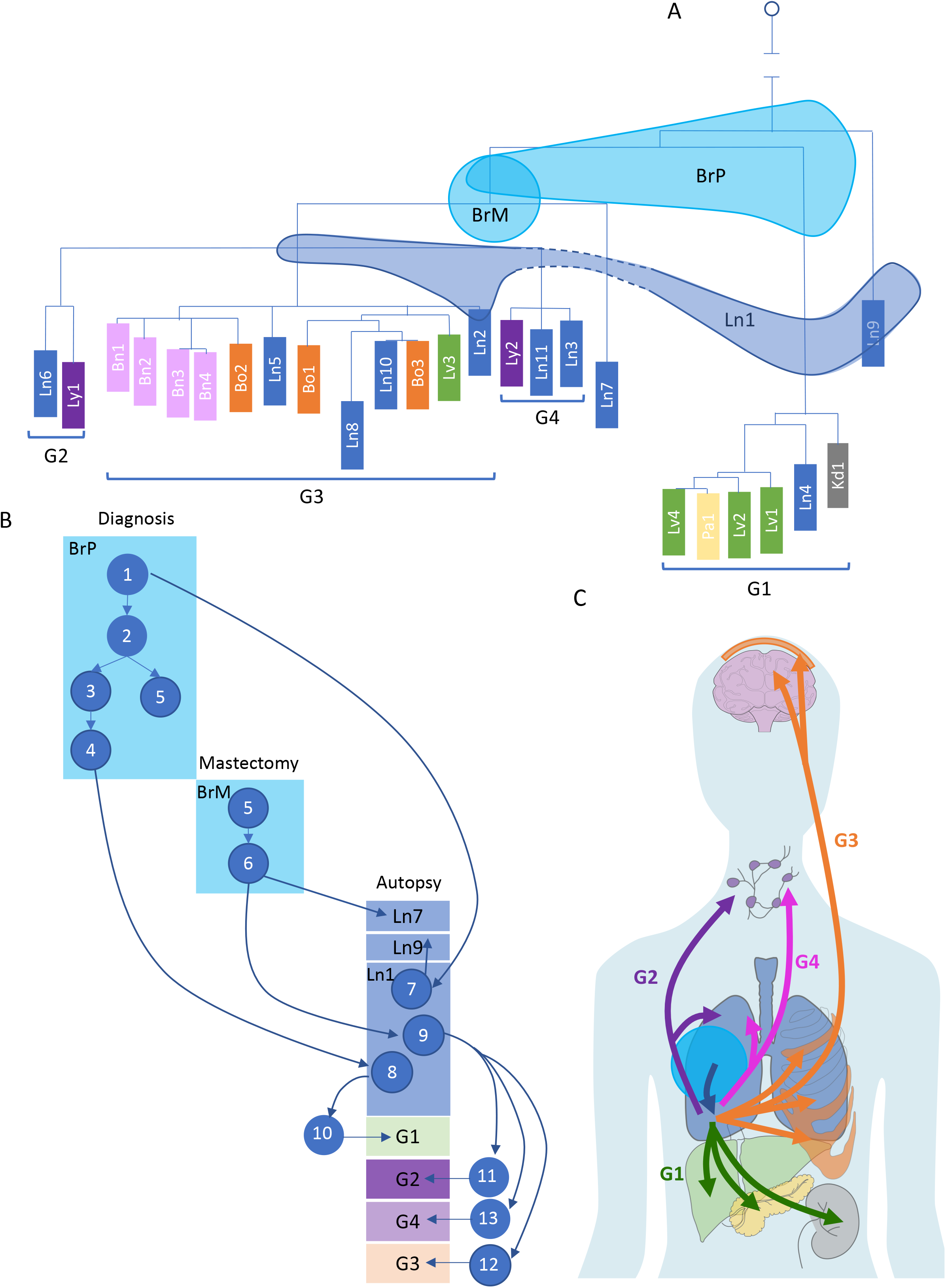
Short variants reveal high-resolution phylogenetic relationship among all tumor biopsies, and early subclonal expansion and migration events. **(A)** Reconstructed phylogenetic tree based on short variants confirms the grouping of the samples and reveals more detailed relationships among samples in groups as well as the placement of Ln7 and Ln9. BrP, BrM and Ln1 shared variants with different samples, indicating that they are mixtures of cell populations. Three shades spanning multiple branches represent sample BrP, BrM and Ln1. **(B)** Reconstructed subclone structures in BrP, BrM, and Ln1 show the heterogeneity of these three samples. The descendant relationship among the subclones suggest the earliest invasion was to lung, and reveal other subclone migrations and expansions from BrP. Each blue circle represents an inferred subclone. Each box represents a sample or a group of samples. **(C)** Overall migration patterns across all tumor sites, with arrow colors corresponding to metastatic waves. After the first invasion from primary breast tumor to lung (blue arrow), four metastatic waves spread the tumors to other organs.

The sample phylogeny derived from chromosome 14 somatic mutations is concordant with the CNV-based grouping, confirming that G1 was seeded by an earlier ancestor, and that the remaining three groups (G2-G4) share a common ancestor (***Supplementary Fig. S4A***). These data also allow placement of two additional lung samples (Ln7 and Ln9) onto the phylogenetic tree. Based on this phylogeny, most of the tumor evolution (i.e. 85%, as measured by the accumulation of somatic mutations, ***Supplementary Fig. S4B***) had already taken place in the primary tumor, before diagnosis. Once the cancer was able to metastasize, however, it rapidly spread, and led to the patient’s death. Three samples (BrP, BrM and Ln1) share mutations with samples across different groups, and therefore cannot be placed concordantly as nodes on the phylogenetic tree. Indeed, these samples span multiple branches of the phylogenetic tree (see colored shade on ***Fig. 3A***), and are likely to be mixtures of cell populations (*i*.*e*. tumor subclones) also present in other samples, a conclusion consistent with the mixed CNV profile observed above for these three samples (***Fig. 2A***). In addition, we refined phylogenetic tree structures in each group using many additional somatic mutation events on copy number-neutral but LOH chromosomes (chromosomes 2, 7, 9, 11, 15, 16, 21 and 22).

### 5. Subclone-level analysis elucidates the main patterns of metastatic evolution in the patient

To understand the composition of these three heterogeneous sites at subclonal resolution, as well as the time ordering and trajectory of the major events that were crucial for this patient’s metastasis, we carried out subclonal analysis based on the mutation allele frequencies of somatic events. In addition to somatic mutations on copy number-invariant chromosome 14, we were able to use many additional somatic mutation events on copy number-neutral but LOH chromosomes (chromosomes 2, 7, 9, 11, 15, 16, 21 and 22), in order to refine group-specific subclone structure. Each subclone in this reconstruction is defined, on average, by 33 mutations (see **Methods**; ***Supplementary Fig. S5-9; Supplementary Table 3-6***). This analysis revealed the complex subclonal composition of the primary and mastectomy breast tumors (BrP and BrM) as well as lung metastasis Ln1, and elucidated the critical role that the subclones present in these key, but heterogeneous samples, played in the metastatic process within our patient (***Fig. 3B***, blue circles labeled with numbers represent the corresponding subclones). Notably, lung metastasis Ln1 was the site of the initial metastatic escape from the breast. This site contains subclone Sc8 that was derived from primary subclone Sc4 (***Supplementary Fig. S5A***). However, neither Sc4 nor any of its descendant subclones are present in the mastectomy (sample BrM), indicating that lung site Ln1 had likely already been colonized before the mastectomy procedure took place (***Supplementary Fig. S5A***). Furthermore, although primary subclone Sc1 is inferred by our subclonal analysis, this ancestral subclone is no longer present at the primary site (BrP) at the time and location of resection (the observed subclone frequency of Sc1 is zero) (***Supplementary Fig. S5D***). This indicates that metastatic site Ln1 was established even earlier, i.e. before the time of the patient’s diagnosis and primary tumor resection. Our analysis also revealed that all four metastatic waves in the patient were seeded by subclones from lung site Ln1: subclone Sc8 gave rise to the first metastatic wave (group G1) (***Supplementary Fig. S5A***); and Sc9 to the three later metastatic waves (G2-G4) (***Supplementary Fig. S5C***). These observations establish lung site Ln1 as a “jumping board” for all subsequent metastatic spread in this patient (***Fig. 3C***).

Previous studies have reported examples of monoclonal and polyclonal seeding in breast cancer patients[7,8,16]. We observed both of these patterns in our patient (***Fig. 4***). As the most striking example of *monoclonal seeding*, multiple subclones from lung site Ln10 seeded as many as nine metastases, primarily in the brain and bones (***Fig. 4; Supplementary Fig. S8***). Subclone Sc17 colonized five distinct sites, including all four metastases in the brain (***Fig. 4; Supplementary Fig. S8B***). The lack of additional mutations across these sites suggest that they were formed within a very short time period (in contrast to Sc18-21, assuming a similar mutation acquisition rate across sites). Conversely, subclones Sc18-Sc21 each seeded a single metastatic site (***Fig. 4; Supplementary Fig. S8B***). These subclones evolved from each other by accumulating mutations gradually and colonizing additional sites in a stepwise manner. This pattern demonstrates that lung site Ln10 acted as a ***subclonal incubator***, in which subclones were able to evolve before colonizing consecutive sites. We also found examples of *polyclonal seeding* (***Fig. 4***, green arrows): subclones Sc1 and Sc4 from the primary tumor colonizing lung site Ln1 (***Supplementary Fig. S5***); and subclones Sc24 and Sc26 from pancreas site Pa1 colonizing liver site Lv4 (***Supplementary Fig. S6***). The important distinction between these two polyclonal seeding events is that the former represents *primary tumor to metastasis* seeding, whereas the latter is a *metastasis to metastasis* event. It is important to note that in this patient, the vast majority of the metastatic colonization events fall into this latter category *i*.*e*. originated from an already metastatic site, rather than directly from the primary tumor, consistent with recent findings in other metastatic cancers[8,13,17]. Finally, *metastatic recolonization* of an existing tumor site has been noted in a cell line engrafted mouse model[18]. We observed such a recolonization event in this patient (***Fig. 4***, red arrow): a subclone (Sc9) and its further evolved descendent (Sc16) are observed at a single site (Ln1). However, subclones Sc12 and Sc14, representing intermediary evolutionary steps between Sc9 and Sc16, are found at a different lung site, Ln2 (***Fig. 4, G3 Ln2; Supplementary Fig. S8A***), and at that site only. The most parsimonious explanation for this observation is that, after evolving at site Ln2, subclone Sc16 invaded, i.e. “recolonized” site Ln1, an already established metastatic site.

**Figure 4.**
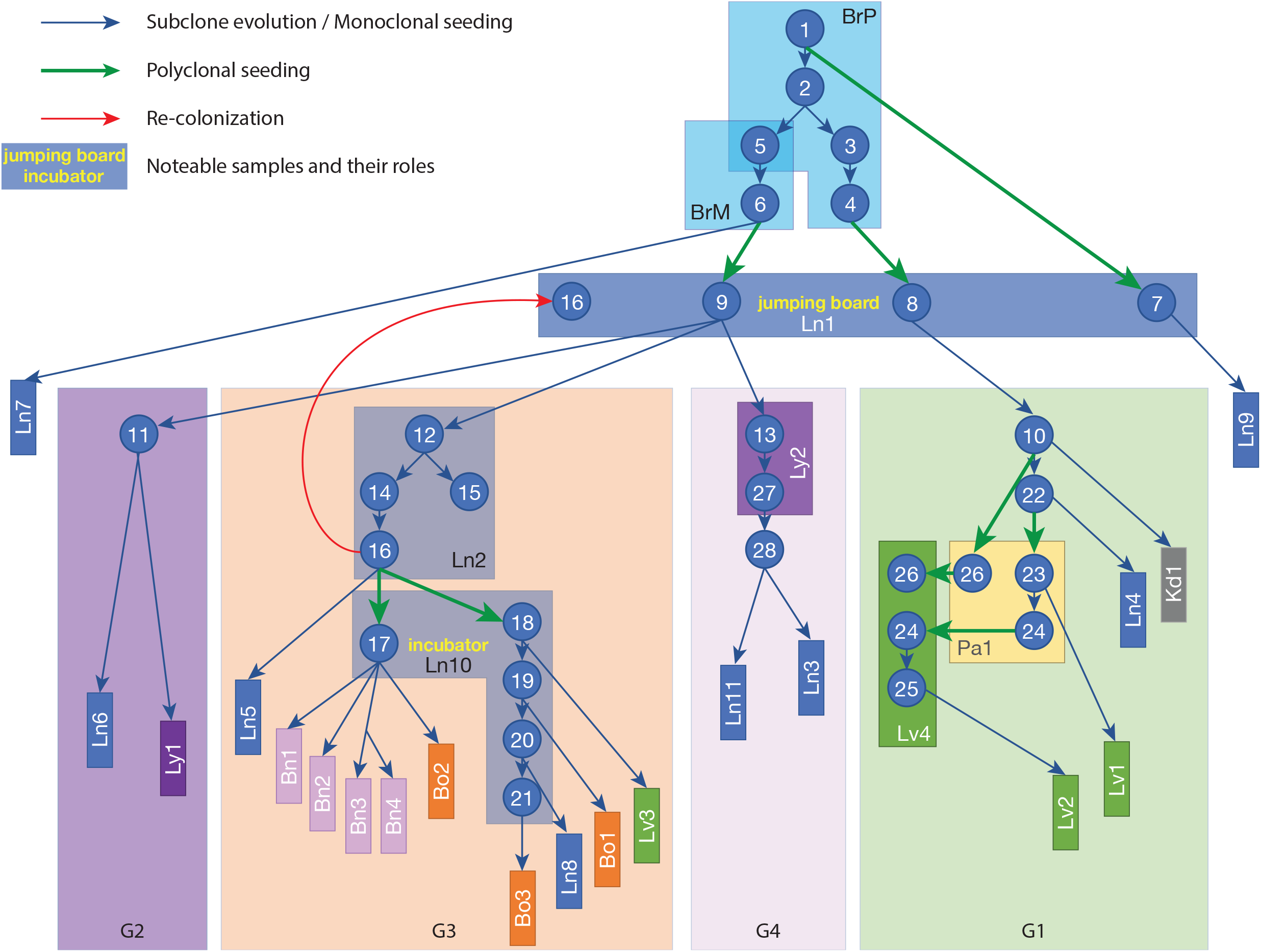
Subclone evolution and migration across all samples. Each blue circle represents an inferred subclone. Each box represents a sample or a group of samples. Samples in the same group are shown in a group box labeled G1-G4. Blue arrows represent subclone evolution or monoclonal seeding. Green arrows represent polyclonal seeding (Sc1, Sc4, and Sc6 seeded Ln1; Sc26 and Sc24 seeded Lv4). Red arrow represents recolonization (Sc16 from Ln2 recolonized Ln1). Ln1 served as the “jumping board” where Sc7 and Sc8, descendants of Sc1 and Sc4 from BrP, and Sc9, descendant of Sc6 from BrM, were all attracted to stay, further evolved, and then colonized other samples. In G3, Ln10 served as an “incubator” where many subclones evolved, and went on to seed other sites.

### 6. Reconstruction of metastatic evolution is robust with respect to unsampled sites

An important question for studies of metastatic cancer evolution is the impact of “missing” *i*.*e*. unsampled metastatic sites on the accuracy of the evolutionary patterns reconstructed from the sampled sites. In this study, 20 of 28 total tumor sites are phylogenetic “leaves” that give rise to no further sites. Were any one of these sites left out of the study, our conclusions would be minimally affected. However, leaving out one of the more complex sites for which our analysis inferred a more critical role would have a larger effect. For example, had we not been able to collect a biopsy at lung site Ln1 (***Supplementary Fig. S10A***), we would not have observed a metastatic recolonization event, or been able to identify the lung as the critical first site of metastatic escape. Without this site, we would have concluded that the primary tumor site (BrP) gave rise directly to the first metastatic wave (G1), and that the mastectomy site (BrM) gave rise to the three later metastatic waves (G2-G4). As a result, substantial insight into the role of the lung in the metastatic spread in this patient would have been lost, although the inferred overall trajectory of metastatic evolution would not have been affected. Unavailability of sample Ln2 would have resulted solely in the loss of our ability to infer the recolonization event (***Supplementary Fig. S10B***). Losing sample Ln10 would have obfuscated the identification of the “incubator effect” in the lung. Unavailability of either sample Lv4 or Pa1 would have resulted in the loss of observation of a polyclonal seeding event (***Supplementary Fig. S10C***). Overall, the loss of most tumor sites would have had localized impact on the conclusions, without distorting the global evolutionary patterns reconstructed by our methods. We note that these considerations also address sampling biases due to *spatial heterogeneity* i.e. subclones at a tumor site potentially missed in the biopsy or autopsy sample.

### 7. Functional annotation of the somatic variants explains the aggressive metastatic disease observed in the patient and provides a window into organ group-specific metastasis

To assess the driver mechanisms for tumorigenesis and metastasis, we interrogated the overall mutation signatures, mutations on well-known oncogenes and tumor suppressors, mutations on genes that have been reported to be involved in metastasis, as well as metastatic sample and group-specific mutations.

We first examined the relative contributions of COSMIC mutational signatures at different stages of tumor evolution (see ***Fig. 5A***). We observed that signatures 1, 3, 5, and 8 were consistently present throughout the tumor’s evolution, suggesting that the mutational processes that caused these signatures were continuously present throughout the disease progression. Signature 1 and 5 are correlated with age of diagnosis and signature 3 and 8 are associated with homologous-recombination deficiency (HRD). Consistent with the previous studies [19,20], TP53-mutated, relatively late diagnosis, TNBC patient had enrichment in signature 3. The dominant signatures during the early evolution of the tumor (i.e. those associated with truncal mutations, present at each tumor site), point overwhelmingly to APOBEC activity (signatures 2 and 13). Strikingly, these signatures were almost completely absent during the later stages of tumor evolution, pointing to cessation/attenuation of APOBEC activity at metastatic sites, in contrast with the general pattern that APOBEC mutational signature increases in prevalence during the course of tumor evolution[12,21,22]. Nevertheless, this phenomenon has been observed in the Yates *et. al*. study, where in one of the studied patients’ tumor, APOBEC activity-related mutational signatures was present early on, but was later turned off[23]. Signature 18 was found to be only present in sample-private mutations (Fig. 5A). Signature 18 was previously reported to be associated with oxidative DNA damage due to reactive oxygen species (ROS)[24,25], which can be induced by gamma-radiation[26]. A recent study[27] also showed that gamma-radiation can induce mutations linked to signature 18. The high prevalence of signature 18 that presented in the late tumor evolution may be due to the radiation therapy this patient received after the mastectomy.

**Figure 5.**
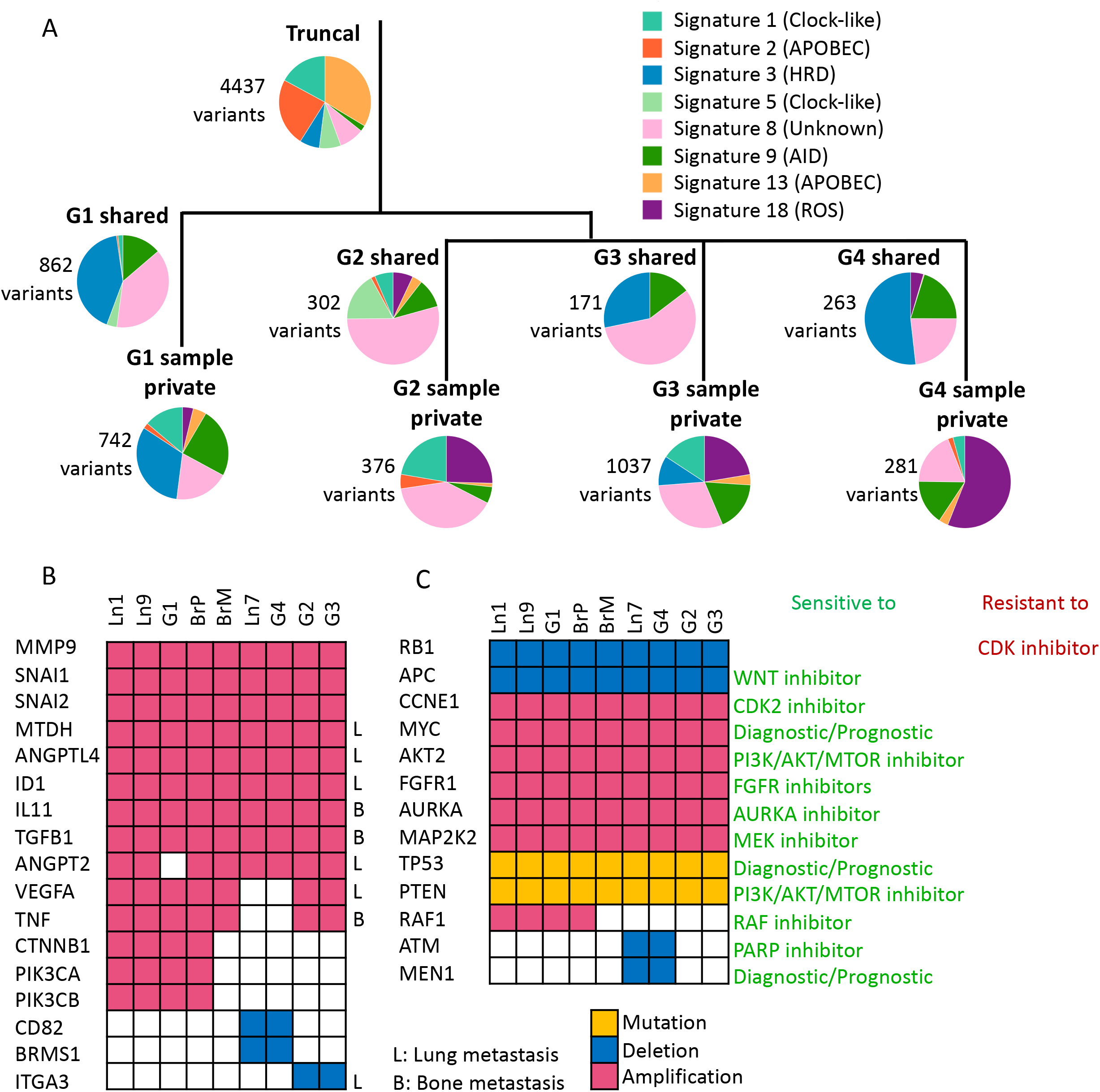
Temporal and spatial distribution of mutational signatures, mutated metastasis-related genes and clinically actionable genes. **(A)** Genome-wide mutational signatures exhibited at different stages of tumor evolution, annotated to the phylogenetic tree. The number of variants is labeled beside each pie chart. HRD, homologous recombination deficiency. ROS, reactive oxygen species. **(B)** Distribution of metastasis related mutations across genes and samples. Genes that are involved in lung metastasis labeled L; genes that are involved in bone metastasis labeled B. **(C)** Distribution and type of potentially clinically informative mutations across genes and samples. Gene mutations are shown in yellow, deletions in blue, and amplifications in red.

Second, we tried to identify driver mutations. A homozygous *TP53* missense (c.517G>C, p.V173L) SNV, a homozygous *PTEN* frameshift (c.676_697delTCCTCCAATTCAGGACCCACAC, p.S226fs), and a homozygous *RB1* deletion occurred in the founding subclone, indicating that these mutations were major primary driver mutations in this patient’s tumor. Then we focused on mutations in 128 genes (***Supplementary Table 7)*** that have been reported as metastasis-related[28–30]. We found that this patient had somatically acquired deletions and amplifications in 17 of these genes (***Fig. 5B)***; eight of the genomic lesions were present in the primary tumor, including the *MTDH, ANGPTL4*, and *ID1* genes associated with lung metastasis[29,31]; and the *IL11* and *TGFB1* genes in which genomic alterations have been associated with bone metastasis[29,32].

Third, we identified mutations in those metastasis-related genes that may contribute to sample-or group-specific tumor cell survival and expansion (***Fig. 5B***). Three of the previously reported metastasis-related genes (*CTNNB1, PIK3CA, PIK3CB*) were somatically altered only in group G1 samples, as well as in the G1-progenitor subclones within samples BrP and Ln1; these genes are therefore candidates for promoting abdominal organ-specific metastasis.

We speculate that some of the nonsynonymous coding mutations that were shared among all samples in the same group, albeit without well-established links to tumorigenesis and metastasis, may also contribute to group-specific metastasis (genes *HSPG2* and *ISM1* are mutated in samples in G1; *IQGAP2* and *RUNX1T1* in G2; *SPINK5* and *FGA* in G3; and *PCDH20* in G4) (***Supplementary Table 8*** and ***Supplementary Fig. S11***). More functional studies are needed, however, to critically evaluate these findings in the context of tumor development.

### 8. Transcriptomic profiling confirms the genomic observation of distinct metastatic waves and identifies group-specific phenotypes

In order to gain insight into the evolution of the patient’s tumor on a molecular / phenotypic level, and thus to complement genomic findings, we collected bulk RNA-seq data from the 26 rapid-autopsy tumor biopsies and 14 surrounding normal tissue samples. First, CNVs were inferred from RNA-seq data by averaging expressions across 101-gene windows and normalized to the expression of 14 normal samples (**Method, *Supplementary Fig. S12A***). 20 tumor samples showed significant copy number changes and recapitulated the presence of large CNVs from the WGS data (major CN events from WGS data were highlighted in dotted boxes in ***Supplementary Fig. S12A, 12C***). The remaining 8 tumor samples showed no copy number changes, which was likely due to heavy normal tissue contamination. Indeed, PCA analysis and unsupervised k-means clustering concordantly showed that the samples without CNV events clustered together with the normal samples (***Supplementary Fig. S12B***), thus we excluded these 8 samples from subsequent analysis. Then, we explored what factors played roles in shaping the transcriptomes of the metastases. Unsupervised clustering on 20 metastatic tumor samples showed that although some samples clustered along their genomic groups (e.g. Ly1 and Ln6, both belonging to G2, were clustered together), and some samples clustered along their host tissue types (e.g. most of lung metastases clustered together; all three brain metastases clustered together), neither genomic grouping nor host tissue types were the sole factor that affected the expression (***Supplementary Fig. S13A***). On one hand, via clustering samples located in the lung and liver respectively because these two organs hosted multiple metastases from different genomic groups, we found that samples in the same genomic group were in fact clustered together (e.g. Ln5, Ln8 and Ln10, which all belong to G3, were more similar with each other than with Ln6 of G2 and Ln3 of G4; Lv1, Lv2 and Lv4, which all belong to G1, were more similar with each other than with Lv3 of G3; ***Supplementary Fig. S13B***,***C***). On the other hand, we clustered the samples that belong to G3, which contained the most diverse host organ types, and found that samples located in the same organ tend to cluster together (***Supplementary Fig. S13D***). This indicates that the environment of the host tissue can shape the transcriptomes of the subclones from the different genomic lineages after these subclones landed and made them adaptive to the respective organs. Next, we compared the gene expression among metastatic samples that belong to different genomic groups. Differential expression (DE) analysis showed that there were 415 genes that were significantly different among the groups (ANODEV test FDR<0.05, logFC>2). Particularly, genes involved in beta1 integrin cell surface interactions (*FGB, FGA*, and *F13A1,VTN, MDK*) were upregulated in G1; genes involved in Wnt signaling network (*FZD8, FZD10, DKK1*) that upregulated in G2; genes involved in integrin family cell surface interactions (*ITGA10, ITGB7*) and in estrogen receptor alpha network (*ESR2* and *GREB1*) were upregulated in G4. Genes that are upregulated in G3 are involved in multiple mechanisms: *RAB40B* can promote tumor cell invasion by regulating trafficking MM2 and MM9 during invadopodia formation[33,34]; *RET* activation can drive signaling through MAPK and PI3K pathways[35]; *ALDH1A1* is a marker for cancer cell stemness[36]; *ST6GALNAC1* encodes the protein in the same family member as *ST6GALNAC5* which has been reported that can mediate infiltration into the brain[37]. Considering the samples in G3 located in multiple tissues, these genes may increase the fitness and invasiveness of samples in G3 to be able to colonize multiple distal organs. This result showed that subclones from different genomic groups probably rely on different survival mechanisms/strategies. Gene set enrichment analysis (GSEA) showed that copy number is the dominant factor in G1 and G3 group (e.g. gene sets NIKOLSKY_BREAST_CANCER_17Q11_Q21_AMPLICON and 17Q21_Q25_AMPLICON were enriched in samples in G3; gene sets 8Q12_Q22_AMPLICON and 8Q23_Q24 were enriched in samples in G1) and consistent with the underlying genetic alterations.

### 9. Delineation of tumor evolution suggests alternative treatment strategies

We identified all somatic mutations (CNVs and short variants) in our tumor samples that impacted a gene with known and clinically targetable mutations in the TARGET database[38,39]. This allowed us to evaluate genetic information that may have impacted the patient’s treatment had it been available to the treating oncologist while the patient was still alive. We found that 11 of 20 (***Supplementary Table 9***) potentially targetable alterations were already present in the primary tumor at diagnosis, and were subsequently retained in all metastatic samples (***Fig. 5C***). These alterations include *RB1* loss, *APC* loss, *MYC* amplification, and *AKT2* amplification; as well as alterations involving genes with high cancer patient population frequencies in the *METABRIC* dataset[40], *i*.*e. FGFR1* amplification (14% patient population frequency) and *AURKA* amplification (6% patient population frequency). These mutations would have been the optimal targets for therapy to impact all metastatic sites in the patient. 5 of the 20 were potentially targetable, sample- or sample group-specific alterations: *MEN1* loss is present only at lung site Ln7 and in G4 samples; *CTNNB1, PIK3CA, PIK3CB* and *RAF1* are amplified only in G1 samples as well as in the samples containing their progenitor subclones (i.e. BrP, Ln1) (***Fig. 5C***). Because these mutations are only present in some but not all metastatic sites, therapies targeting these genes are unlikely to have been effective. This observation underlines the necessity of comprehensive monitoring of metastatic sites for effective therapeutic intervention.

## Discussion

The 46 biopsy / autopsy samples in this dataset, the largest number from a single patient to date, allowed us to track the evolution of this metastatic tumor genome at subclonal resolution, as it spread from the breast to seven additional organs (***Fig. 6***). Reconstruction of the evolutionary trajectory of the tumor revealed four distinct waves of metastatic colonization, targeting well-delineated groups of organs in the patient.

**Figure 6.**
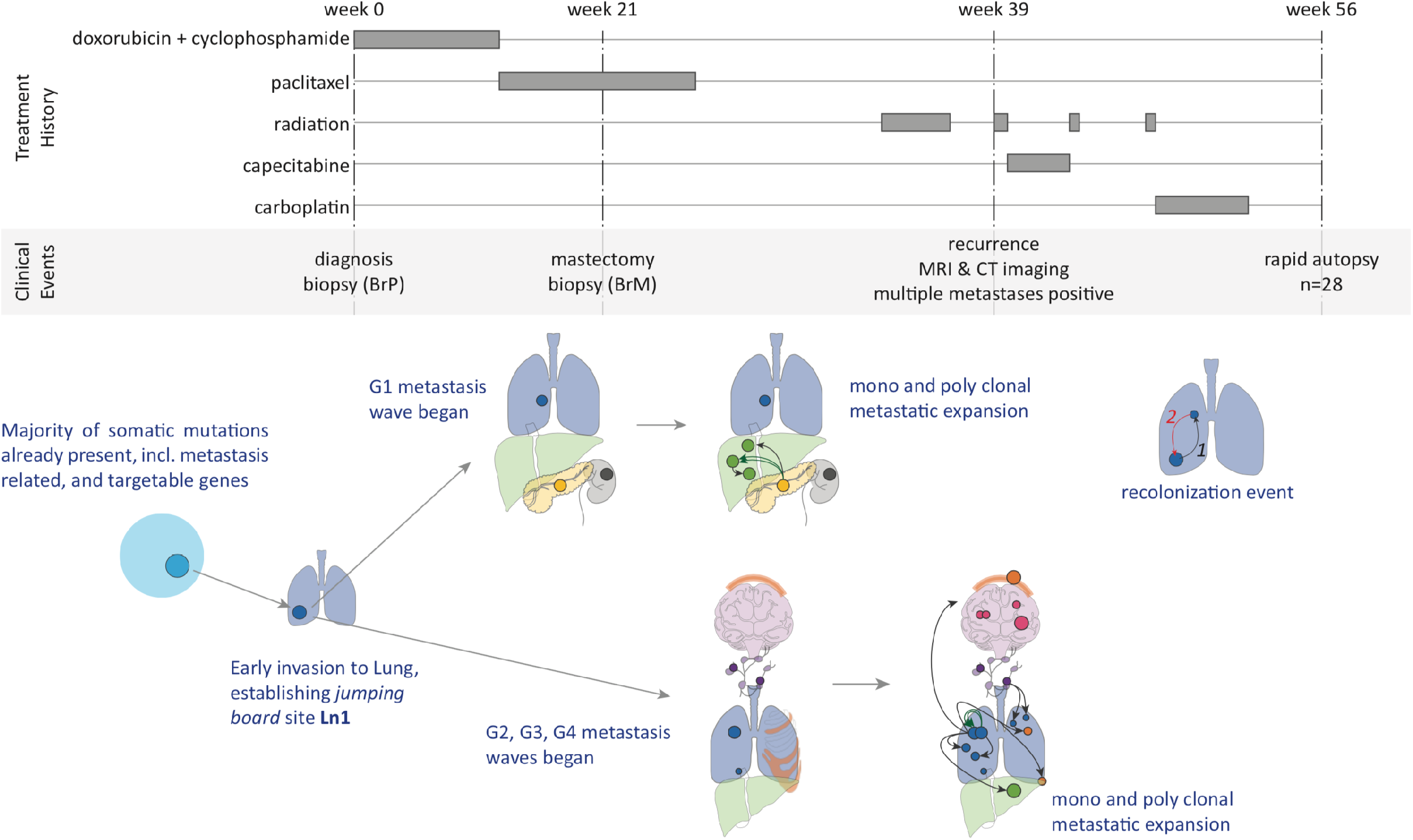
Summary of tumor evolution and metastasis progression in correlation with treatment history and clinical events.

Our study reveals the existence of “metastatic incubators” where subclones that initially colonized the sites further evolved into founding clones of additional sites. There are two independent lines of evidence for our conclusion that these subclones evolved at metastatic sites, rather than that they were already present in the primary. First, we observed the evolutionary intermediate clones at lung sites Ln1 (Sc8, Sc9), Ln2 (Sc14-Sc16), Ln10 (Sc17-Sc21), whereas we did not find any of these intermediate subclones in the breast samples (i.e. in BrP or BrM), or any other sample in our collection. This suggests *in situ* evolution at, rather than migration from the primary to, those sites. We note that the same argument holds true for other organ sites where evolutionary intermediaries were observed, i.e. liver site Lv4 (Sc24), lymph node site Ly2 (Sc27), and pancreas site Pa1 (Sc24). Second, the evolutionary intermediate subclones are not defined by one or two, but by multiple somatic variants (***Supplementary Table 3***). The absence of these variants in BrP and BrM suggests that the intermediate subclones were absent from the breast samples. Previous breast cancer studies [6,8,9] observed mutations not found in the primary tumor but shared between multiple metastatic sites, and suggest the presence of an already metastatic, common ancestor subclone seeding those sites. Our study provides direct, high-resolution evidence for the presence of such common ancestors clones, and elucidates the great extent of subclonal evolution that occurred at these sites. Our study highlights the critical role of the lung in the metastatic process of our patient. The lung was not only the first site of metastatic escape, but also an organ with multiple metastatic incubator sites. A larger TNBC metastasis cohort with similar genomic resolution is needed to validate that the lung is a metastatic incubator hotspot, or to identify other incubator “hotspot” organs.

The presence of somatic alterations in genes associated with lung metastasis in the primary tumor is consistent with our subclonal migration results (see Sections 4 and 5) indicating that the lung was the first organ of metastasis. The scenarios for distal organs are more complicated. On the one hand, the presence of somatically altered genes associated with bone metastasis in the primary tumor signals that the tumor’s ability to metastasize to the bone may, in whole or in part, be already there in the primary tumor. On the other hand, our subclone analysis revealed that all distal organ metastasis sites outside the lung had originated not directly from the primary site, but from lung metastasis sites by new subclones harboring additional mutations. Taken together, these observations suggest that the rampant metastatic invasion was the compound result of early mutations in the primary tumor and later evolution at metastatic sites in the lung.

Once metastasis was established, resection of the primary (breast) site was ineffective against the progression of this aggressive disease. This reinforces the Fisher hypothesis that breast cancer is a systemic disease in which metastasis occurs before diagnosis[41]. Although many potentially targetable mutations were already present at the primary site, and therefore at all tumor sites in the patient, others were only present at a subset of metastatic sites. This highlights the need for more comprehensive sampling of metastatic sites: indeed, targeted treatment strategies identified from assays of the primary site, or only one or two additional metastatic sites, may fail to achieve the desired therapeutic outcome[42]. Therefore, a more comprehensive survey of metastases (or liquid biopsy-based approaches[6,43] if direct sampling of additional metastatic sites is not practicable) may be necessary for guiding more successful clinical interventions.

This study lays the methodological foundations for tracking a patient’s metastases at subclonal resolution. With additional, similarly comprehensive datasets in hand, it will be possible to assess the generality of these findings, and to establish how the specific evolutionary patterns observed in a patient’s metastatic tumor evolution can, in the future, inform more effective personalized treatment.

## Conclusions

This study reports on a one-of-a-kind triple negative breast cancer (TNBC) metastatic tumor dataset. This unique dataset allowed us to track the patient’s tumor across metastatic colonization in eight distinct organs, at subclonal resolution. The most striking finding of this study is the observation of “metastatic incubators”, where substantial subclonal evolution preceded, and may have been necessary for, further metastatic colonization. We found that the lung played a crucial role in metastatic tumor spread in our patient, a finding that must be examined in larger patient cohorts. Our results showcase the level of detail achievable for reconstructing metastatic subclonal evolution when sampling a large number of metastatic sites, and combining this with deep genome sequence-based subclonal analysis.

## METHODS

The workflow for this study is shown in ***Supplementary Fig. S14***.

### Sample collection

The study was reviewed and approved by the human subjects IRB of the University of Utah. Informed consent in accordance with the Declaration of Helsinki was obtained from the patient. We collected in total 44 autopsy samples including two skin normal tissues, 28 tumor samples and 14 adjacent/distal normal tissues from the tumor (see detailed description in ***Supplementary Table 1***). All metastatic samples represent individual tumors except sample Bn3 and Bn4, and Ln2 and Ln3, which were from different parts of the same brain tumor, and the same lung tumor, respectively. For all autopsy samples, frozen sections were reviewed by pathologists to confirm the tumor type and presence, and to quantify necrosis levels. All autopsy samples were stored in RNAlater at -80C until DNA and/or RNA isolation. FFPE samples for primary tumor biopsy and mastectomy biopsy were also available for this study.

### Sample process

DNA from FFPE samples were isolated using the Qiagen QIAamp DNA FFPE Tissue Kit. DNA from all 28 autopsy tumor samples and 2 skin normal tissues were isolated using Qiagen’s QIAamp DNA Micro Kit. RNA from all 28 autopsy tumors and adjacent/distal normal samples was extracted by using Qiagen RNeasy Micro/Mini Kit.

### WGS analysis including somatic SNV and INDEL calling, CNV calling, LOH calling, structural variant calling, and translocation calling

Primary and mastectomy samples were subjected to 45X WGS at the Huntsman Cancer Institute’s High Throughput Genomics Core Facility using the Illumina TruSeq. Metastatic tumors and skin biopsy samples were subjected to 60X WGS at the McDonnell Genome Institute at Washington University using NantOmics. Samples sequenced at Washington University were provided as aligned BAM files. Primary and mastectomy WGS sequencing data were aligned using an identical pipeline to the one used at the McDonnel Genome Institute to the same GRCh37-lite reference genome (ftp://ftp.ncbi.nih.gov/genbank/genomes/Eukaryotes/vertebrates_mammals/Homo_sapiens/GRCh37/special_requests/GRCh37-lite.fa.gz) using BWA-MEM 0.7.15-r1140; Freebayes 0.9.21 was used to identify SNV and INDEL variants called jointly over all samples using the following command line parameters:

- --allele-balance-priors-off
- --report-genotype-likelihood-max
- --genotype-qualities
- --pooled-discrete
- --pooled-continuous

The variants produced by Freebayes were then subjected to quality filtering, including criteria as follows:

- variant quality > 30
- per-sample sequencing depth > 15
- intersecting with 1000G genome accessibility mask
- inverse-intersecting with low complexity region mask of GRCh37d5
- filtering out multi-allelic variant sites

Somatic variants were identified when the VAF was below 0.1 or the alternate allele count was less than five in both normal skin samples. To ensure that differences in sequencing provider and depths did not affect variant detection, we showed that the number of somatic variants detected in BrP and BrM (45X, at Huntsman Cancer Institute) were similar to other samples (60X, at Washington University). We detected in total 20,012 somatic variants across all samples. On average we detected 10,648 ± 493 (standard deviation) variants per metastatic sample, 10,352 variants in BrP and 10,121 variants in BrM (***Supplementary Fig. S1***). The same trend can be observed for somatic variants on chromosome 14 and chromosomes 2,7,9,11,14,15,16,21,22 (***Supplementary Fig. S1***). SNVs and INDELs were annotated by SnpEFF 4.2.

FACETS[15] was used to identify copy number variants and LOH events. All copy number calls were manually curated. Structural variants and translocations were identified using the reference free variant detection algorithm RUFUS[44] and Lumpy[45] followed by visual inspection in IGV[46,47].

### Allele specific CNV/LOH call in multiple samples

Heterozygosity and copy number for each sample were derived using FACETS. Allele specific copy number changes were not generated by FACETS, but were separately inferred using inherited variants falling in somatic CNV regions. By comparing the AF of these variants between samples, we were able to identify the allele specific copy number changes. For example, the AF of inherited variants on chromosome 3 in pure tumor samples with copy number neutral LOH chromosome 3 is either 1 or 0, whereas the AF of these variants in pure tumor samples with copy number three and both alleles would be 0.33 and 0.67. A scatter plot of AF of these variants between two samples reveals which chromosome is amplified in the copy number amplified sample, as well as whether the amplified chromosome is the same as the ones in the copy number neutral, LOH sample. ***Supplementary Fig. S15A*** shows the AF of inherited exonic variants on chromosome 3 between Pa1 and Bn2. Variants in red circles represent the homozygous variants in both samples. The AF of variants in yellow circles indicate that the amplified allele in Pa1 became the only allele remained in Bn2, at copy number 2. The AF of variants in green circles indicate that the unamplified allele in Pa1 was lost in Bn2. This method enabled us to establish whether samples with the same copy number and LOH are the same events. In addition, this method provides higher resolution information such as allele specific structural variants (including translocation) (***Supplementary Fig. S15B***).

### Phylogenetic tree construction

We used somatic short variants on chromosome 14 (which harbored no copy number or LOH events (except for Ln9)) to construct the phylogenetic tree across all samples and additional somatic mutations on copy number-neutral but LOH chromosomes (chromosomes 2, 7, 9, 11, 15, 16, 21 and 22) to refine the phylogenetic tree in each group. We encoded the state of a known somatic variant locus in a sample as a binary value, where 1 indicates the mutation is present (AF>0.1) and 0 that the mutation is absent (AF<0.1). We used two methods to construct phylogenetic tree. First, we used the UPGMA clustering method based on the hamming distance matrix calculated between samples (***Supplementary Fig. S16***). Samples in the same group were confirmed to cluster. However, this method does not consider the constraint that samples sharing the same variants should share an evolutionary lineage. Therefore secondly, we developed a method that would incorporate this constraint while simultaneously making the assumption that 1) all cancer cells are descendants of a single founding clone (i.e. normal cell); and 2) all mutations satisfy the infinite sites assumption that the chance the same mutation occurs independently in different cells, as well as variants reverting back to the wild type, is extremely low. Therefore, we can describe our problem as a perfect phylogeny problem[48] with complete and cladistic characters which are the states of mutations. For each variant, a binary vector v_i_^j^ is calculated where v_i_^j^ = 1 if variant i is found in sample j, or 0 otherwise. Variants with the same binary vectors are clustered together, which means that they occur in the same clone, albeit the clone can be found in multiple samples. The evolution ordering between any two variants can be established by comparing their binary vectors. If a variant i1 occurred in the clone that already contained i2, for all samples j, either of the following two conditions must hold true: 1) v_i1_^j^ = 1 and v_i2_^j^ = 1 when j contains the descendant clone; or 2) v_i1_^j^ = 1 and v_i2_^j^ = 0 when j contains the ancestral clone before variant i1 occurred. Although Ln9 had acquired an additional chromosome 14, no variants were lost in this process. Thus, this method can still apply to Ln9. The results from the second method showed that Ln9 was the first sample to branch out and the rest of samples had a common ancestor (***Fig. 3A***).

### Subclonal analysis with SubcloneSeeker

We used AFs of somatic variants on chromosome 14 to reconstruct subclone structure and estimate cell prevalence of each subclone of BrP, BrM, Ln1, Ln7 and Ln9. With the exception of Ln9, all samples have CN normal chromosome 14, therefore AF can be used to accurately estimate the cell prevalence. For subclone structure of samples in each group, in addition to variants on chromosome 14, we also used group specific variants that are absent in BrP and BrM on the CN neutral chromosomes containing LOH (chromosomes 2, 7, 9, 11, 15, 16, 21 and 22) events shared by all samples in G1-G4 as well as BrP and BrM. Since these variants occurred after the LOH event chronologically, they are most likely to be heterozygous, and can be used for subclone analysis within a group. Because all samples in G4 had chr11p15-q25 deletion, we can also accurately estimate CP from G4 specific variants in this region. Thus, these variants were also used for subclone analysis in G4.

For subclone analysis, we clustered variants with the same level of AF in all samples to a cluster (C1-C28 in ***Supplementary Fig. S5-9***). We used 0.05 as the allele frequency cutoff for positive somatic variant detection. The ancestral relationship between two subclones satisfy: 1) variants in the ancestral clone have larger AF than variants in the descendant clone in one sample; 2) variants in the descendant clone were inherited from the ancestral clone; 3) variants that have ∼0.5 AF are in the founding clone of a sample. SubcloneSeeker was used for constructing subclone structures and estimated CP for each subclone for individual samples[49].

### Identification of monoclonal and polyclonal seeding

A clone presented at a less than 100% cell prevalence in one sample, and then at 100% in another sample, signifies that this subclone emerged in the former sample, and seeded the latter, which can be characterized as a monoclonal seeding event. However, if a clone had a low cellular prevalence in both samples, it’s likely to be the result of a polyclonal seeding event, in which two or more subclones in one sample traveled together or separately and seeded the other one.

### Mutational signature analysis

We assessed the dynamics of mutational process over time by analyzing somatic mutations patterns attributed to the branches of the phylogenetic tree, including truncal mutations, mutations shared by samples in G1, G2, G3 and G4 respectively, as well as sample private mutations aggregated by their grouping identity. We applied MutationalPatterns[50] to our dataset. Briefly, after de novo extraction of mutational signatures from the mutation count matrix, the contribution of COSMIC mutational signatures (https://cancer.sanger.ac.uk/cosmic/signatures_v2) to the mutational profile were quantified.

### RNA-seq and data processing

RNA-seq was performed using rRNA depletion-based library preparation followed by paired-end Illumina HiSeq sequencing. We obtained RNA-seq data from 42 specimens throughout the patient’s body at autopsy, including 26 grossly tumor (which also had DNA samples) and 16 surrounding normal samples. RNA-seq data were processed with Rsubread[51,52] v1.16.1. We aligned the reads to GRCh37 and used only uniquely mapped reads and the Hamming distance to break ties. The maximum indels allowed per alignment was 5. Gene-level expression values were processed to transcript per million mapped reads (TPM). We used the featureCounts function in Rsubread for reads counting. We used the built-in annotation file which includes the exon annotation information from NCBI Build 37.2 and Entrez gene identifier.

### RNA-seq-based copy number inference

To infer approximate copy number events from RNA-seq data, we used an approach reported previously for single-cell RNA-seq copy number inference[53], which we had also used previously[54]. This approach relies on normalization and calculation of 101-gene window expression averages, followed by normalization to samples with little or no tumor purity.

### Differential expression (DE) analysis

We used a R workflow package, “RnaSeqGeneEdgeRQL”[55] for normalization, and downstream DE and pathway analysis. Specifically, we normalized data by using calcNormFactors function which applied the trimmed mean of M values (TMM) approach. Next, the DE analysis was performed on samples in four groups (G1-G4) by using EdgeR which implemented empirical Bayes methods that permits the estimation of gene-specific biological variations. We made pairwise comparisons between all four groups and performed one-way analysis of deviance (ANODEV) for each gene. FDR<0.05 was used for significance cutoff. Significantly expressed genes in each group were annotated in terms of higher order biological processes or molecular pathways by using NCI-Nature pathway database in Enrichr[56]. Finally, we performed gene set enrichment analysis (GSEA) using the C2 curatd signatures from MSigDB (including 5,637 signatures). RnaSeqGeneEdgeRQL package incorporates the Correlation Adjusted MEan RAnk gene set test (CAMERA)[57] method for the enrichment analysis.

### Validation of variants in RNA-seq data

Somatic SNVs and INDELs identified by Freebayes from WGS data were validated by RNA-seq data. We randomly picked two samples in each group and Ln7 (total nine samples) with high tumor purity in RNA-seq data for this validation. For any given tumor sample, only somatic short variants that have greater than 0.1 genomic VAF, and have a read depth of at least ten in the paired RNA-seq were considered. We use the following workflow to validate the variants:

- If a variant is present in paired RNA-seq data i.e. having RNA-seq reads containing the variant allele, it is considered “validated”
- If a variant is not found in the paired RNA-seq data, but found in the RNA-seq data of other tumor samples genomically determined to also contain the same variant, it is then considered as “validated in other samples”.
- If a variant cannot be validated by either of the mentioned steps, we consider the following possibilities:
  - Variant dropout in RNA seq data due to sampling: It is reasonable to consider that a variant with low WGS VAF (e.g. 0.1) and low RNA-seq coverage (e.g. 10X) may not be sampled by RNA-seq according to binomial distribution (in the example case, the possibility of sampling 0 alternate allele containing reads, or P_0_, is 0.35). We skip such variants with P_0_ > 0.05. Note that less than five variants were in this category in each sample.
  - Variant allele not expressed: this can be the result of unbalanced expression between alleles or false positive in genomic variant calling.

More than 90% variants can be either “validated” or “validated in other samples” (***Supplementary Fig. S17***).

## Code availability

Custom scripts used in this study can be found at https://github.com/xiaomengh/tumor-evo-rapid-autopsy.

## Data availability

Accession code will be available before publishing.

## Acknowledgements

We thank the patient and her family for generously participating in and supporting this research. We thank Kathy Raven from Raven Forensic Pathology & Autopsy Services for conducting a rapid autopsy. X.H., Y.Q., and G.T.M. were supported by U24CA209999 to G.T.M. S.W.B., A.H.B., A.L.C., G.S., and J.A.M. were supported by U54CA209978. A.L.C. was supported by P30CA042014 from NCI. S.W.B. was supported by NLM training grant 5T15LM007124. We thank the University of Utah and the College of Pharmacy for supporting the sequencing of the samples. Research reported in this publication utilized the High-Throughput Genomics and Bioinformatic Analysis Shared Resource at Huntsman Cancer Institute at the University of Utah and was supported by the NCI of the NIH under Award Number P30CA042014. The support and resources from the Center for High Performance Computing at the University of Utah are gratefully acknowledged. The computational resources used were partially funded by the NIH Shared Instrumentation Grant 1S10OD021644-01A1. The content is solely the responsibility of the authors and does not necessarily represent the official views of the National Cancer Institute or the National Institutes of Health.

## Author Contributions

X.H. and Y.Q. jointly contributed to study design, coordination, genomic and transcriptomic data processing and analysis, variant identification, subclone analysis, sample phylogenetic analysis, subclonal migration pattern determination, and manuscript writing. S.W.B. contributed to study design, coordination, sample collection, sample processing for DNA and RNA sequencing, RNA-seq analysis, TCGA subtype identification, and manuscript writing. R.F. and E.D.-K. contributed to pathology analysis of cancer specimens and assessment writing. A.F. contributed to structural variant identification. J.A.M. and G.S. contributed to DNA and RNA isolation. D.J. and W.E.J. helped with RNA-seq analysis. A.L.C. contributed to imaging reviewing, clinical oversight, and manuscript writing. A.H.B. and G.T.M. conceived the project, contributed to project coordination. G.T.M. contributed to manuscript writing. All authors contributed to manuscript refinement.

## Notes

We declare that we have no conflict of interest.

### Competing Interest Statement

The authors have declared no competing interest.

