## Supplemental Figures for "Novel temporal and spatial patterns of metastatic colonization from rapid-autopsy tumor biopsies": Supplental Figures and legends_preprint.pdf

Supplementary Figure S1

A

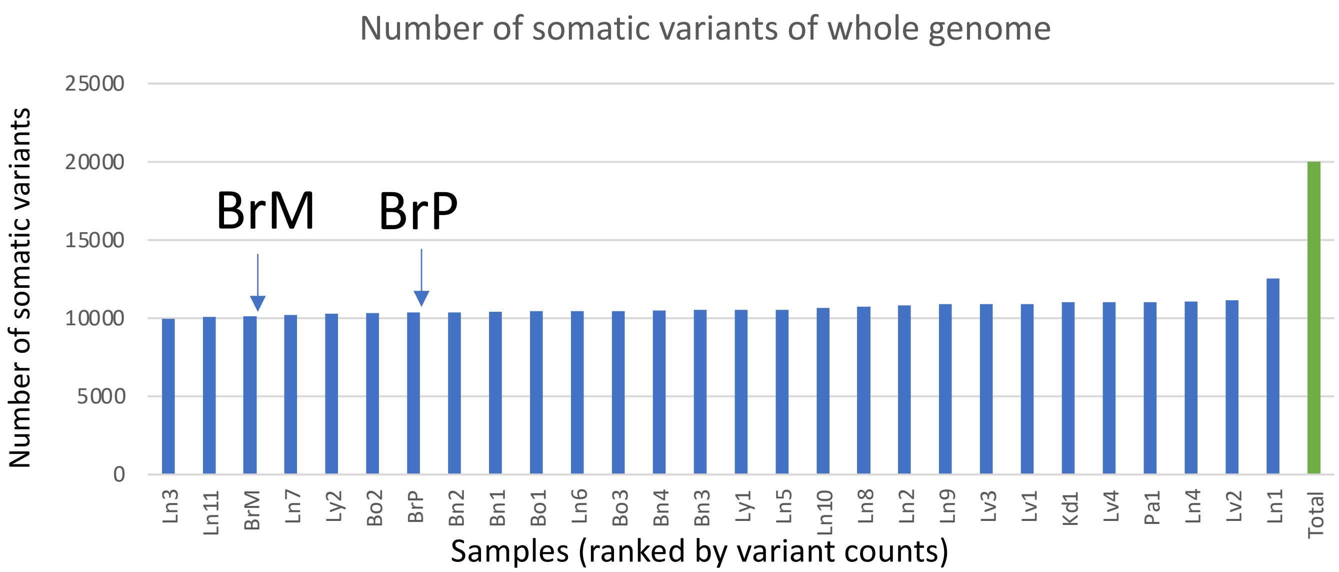


B

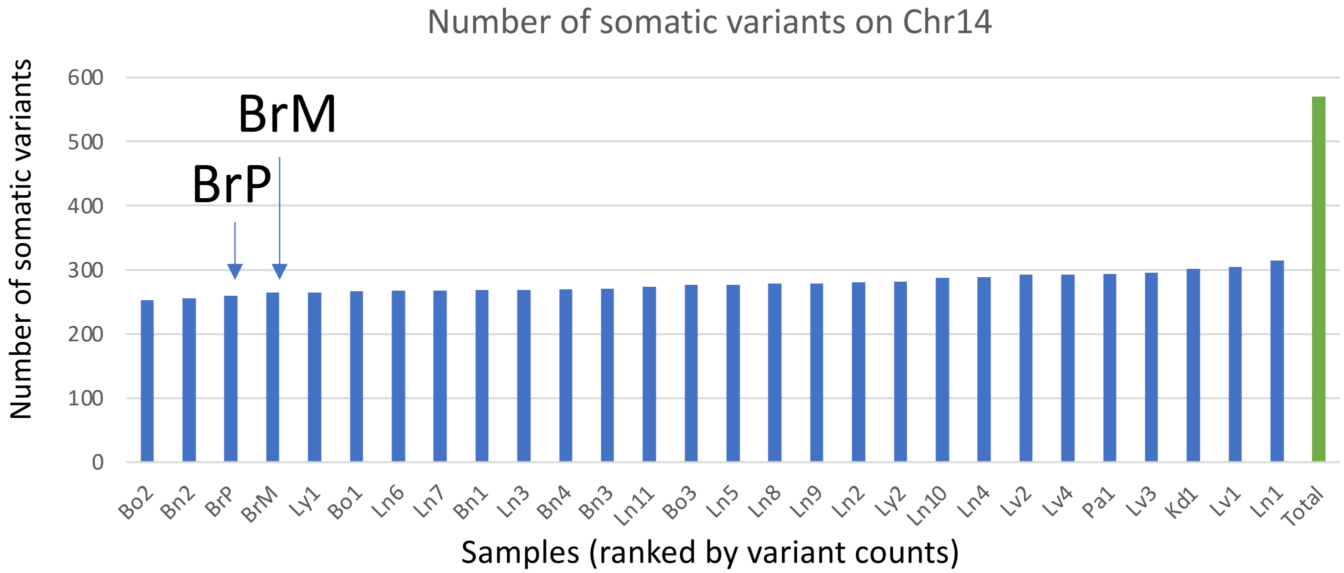


C

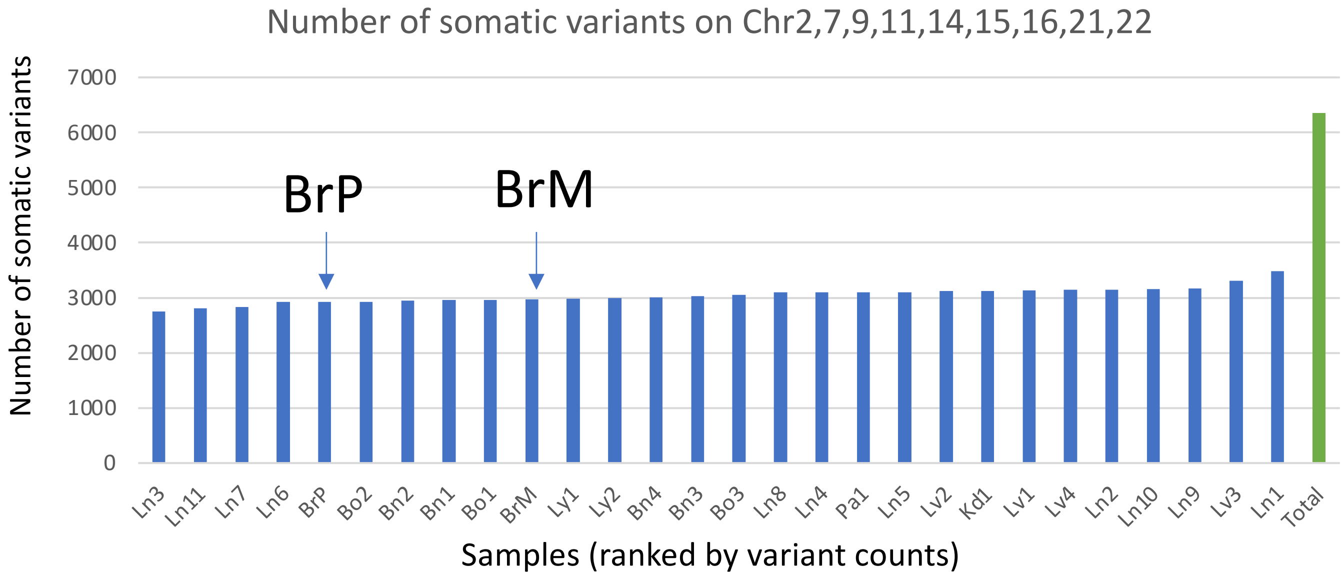

**Supplementary Figure S1. Somatic mutation counts across all tumor samples. (A)** Whole genome somatic mutation counts for all tumor samples. **(B)** Somatic mutation counts on chromosome 14. **(C)** Somatic mutation counts on chromosome 2,7,9,11,14,15,16,21,22. X-axis: samples ranked by variant counts from smallest to largest. Y-axis: the number of somatic mutations. Green bars represent the total number of mutations. BrP and BrM, which were sequenced at 45X coverage, had similar mutation counts as other rapid-autopsy samples sequenced at 60X coverage.

Supplementary Figure S2

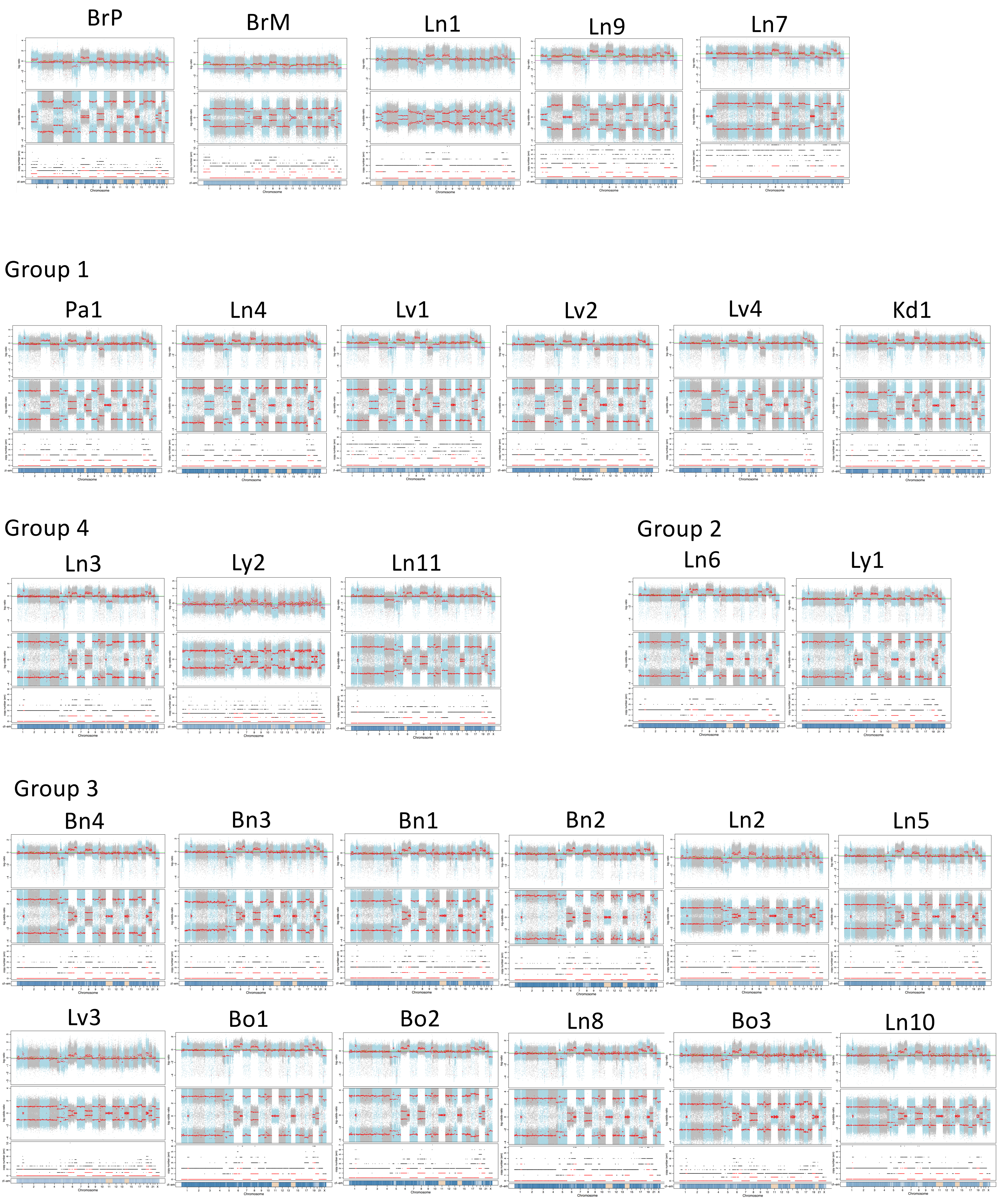

**Supplementary Figure S2. CNV profiles of all samples generated using FACETS.** For each sample, top panel is the log2 ratio of tumor copy number to normal copy number. The middle panel is allele-specific log-odds-ratio with chromosomes alternating in blue and grey. The bottom panel is the corresponding integer (total, minor) copy number calls by FACETS.

### Supplementary Figure S3

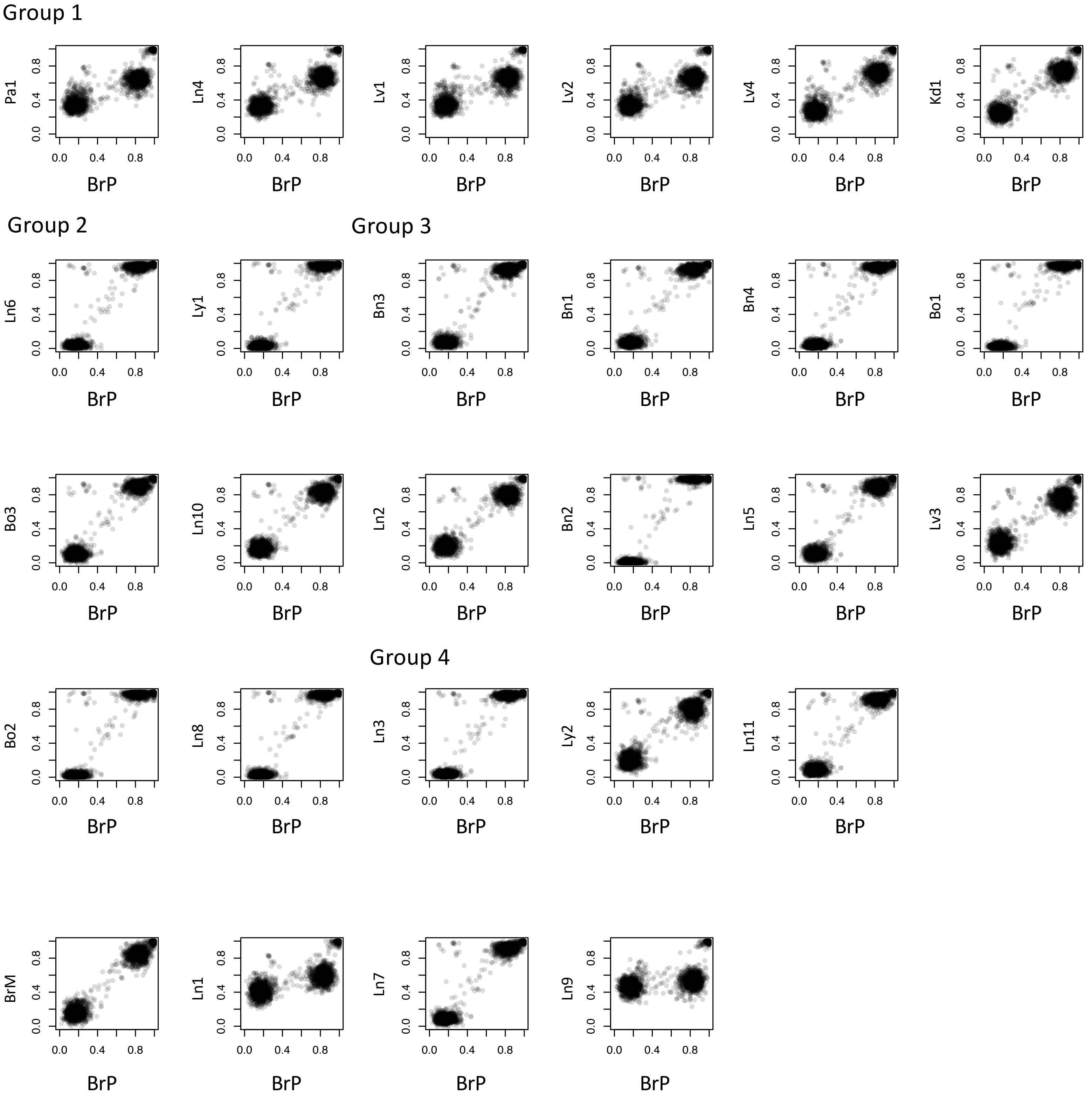

**Supplementary Figure S3. Scatter plot of allele frequencies of inherited variants on chromosome 3 between BrP and other biopsies,** to verify that the amplified allele in G1 was also the amplified allele in BrP and the unamplified allele was lost in G2, G3 and G4.

Supplementary Figure S4

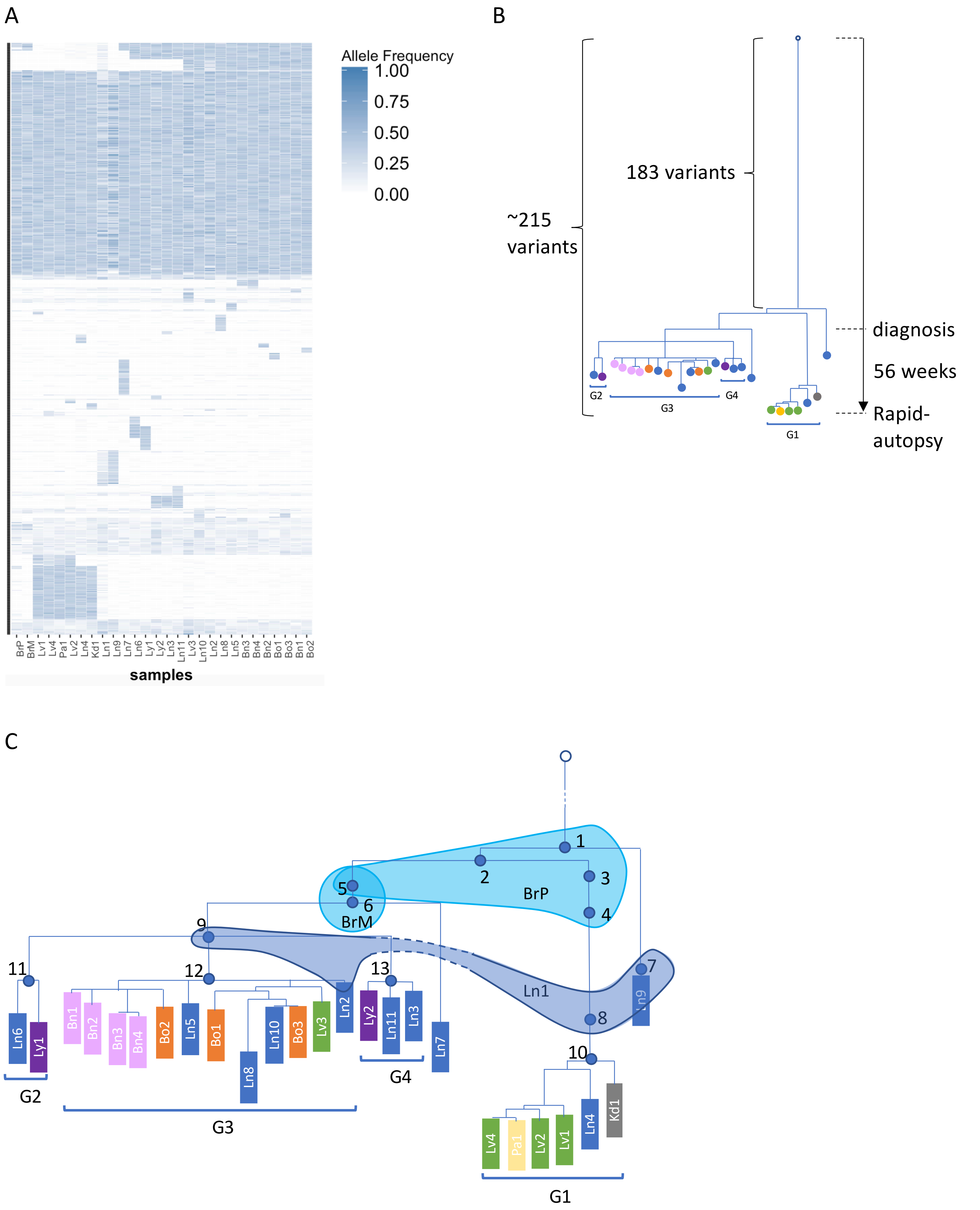

**Supplementary Figure S4. Construct phylogenetic tree based on somatic short variants.** **(A)** Allele frequencies of somatic variants on chromosome 14. Each row is a variant and each column is a sample. **(B)** Phylogenetic tree constructed by a perfect phylogeny algorithm using somatic variants on chromosome 14 considering shared and unique variants among samples. The sample order from the left to right is the same as in (C). The branch length was proportional to newly acquired mutations. There are 183 somatic variants before the first bifurcation whereas on average ~215 somatic short variants on each evolution path. The corresponding time scale over disease progression is shown on the right. **(C)** Revised phylogenetic tree for samples in groups by incorporating heterozygous short variants from chromosomes 2, 7, 9, 11, 14, 15, 16, 21 and 22 showing that LOH events happened in BrP. We placed the inferred subclones (blue dots labeled number 1-13) in ***Figure 3B*** on this phylogenetic tree as their position on the evolution process. Three shades of blue containing different ancestral subclones (labeled BrP, BrM and Ln1) are the highly heterogeneous samples.

Supplementary Figure S5

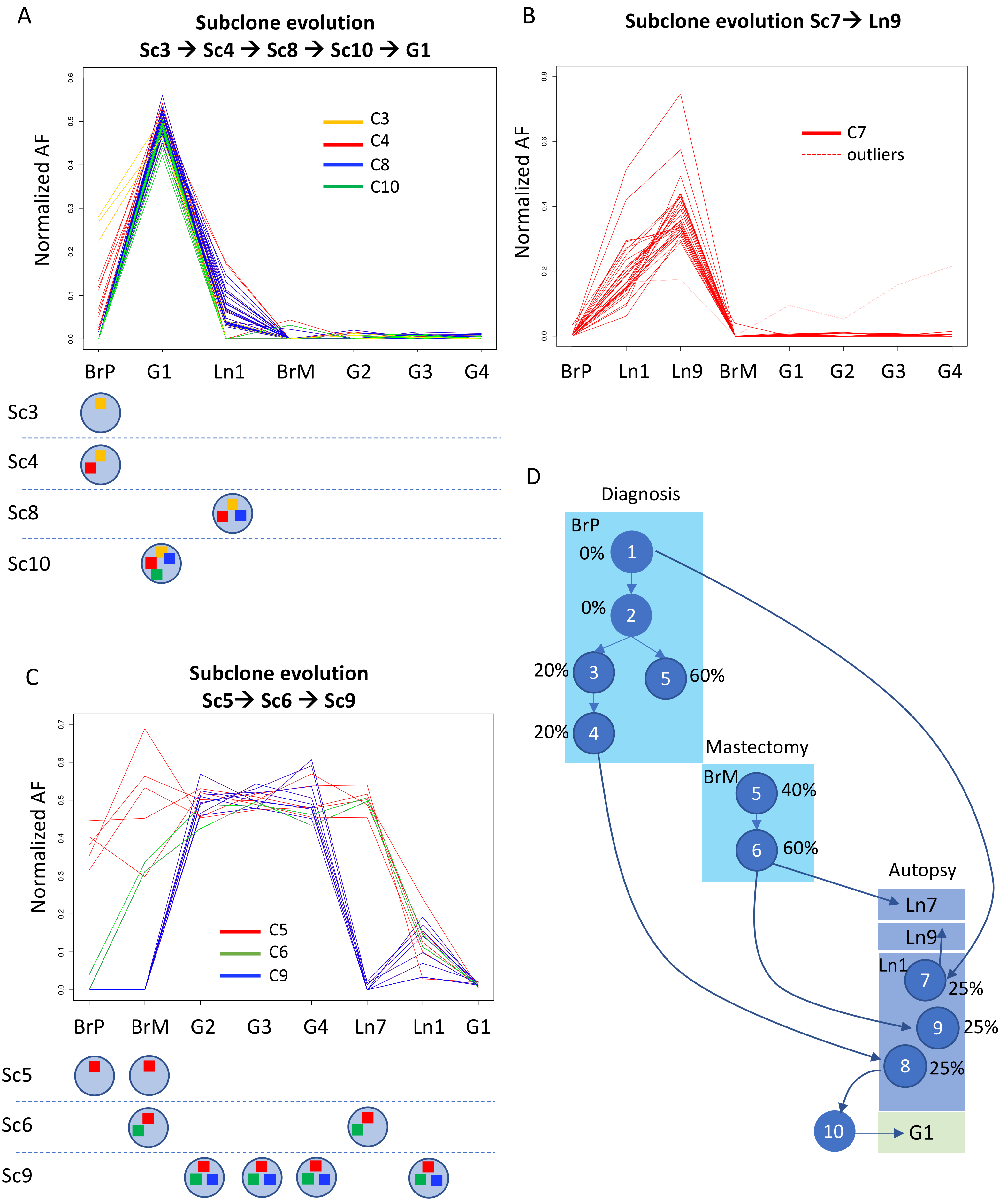

**Supplementary Figure S5. Early subclone evolution and expansion in breast tumors and the first invaded lung sites. (A)** Subclone evolution Sc3->Sc4->Sc8->Sc10->G1. The top panel is normalized AF of short variants on chromosome 14 clustered into C3, C4, C8, and C10 depicted in yellow, red, blue, and green respectively. The bottom panel is a schematic plot of which subclone contains which variant clusters and which sample contains which subclones. Each row is a subclone. A circle represents a subclone and a colored square inside of the circle represents the corresponding variant cluster in the top panel. For example, Sc3 only contains variants in cluster C3, whereas Sc10 contains variants in C3, C4, C8, and C10. A sample contains subclones below this sample. BrP contains Sc3 and its descendant Sc4. A further evolved subclone Sc8 from Sc4 went to Ln1 and eventually became the common seeder Sc10 for samples in G1. None of these subclones are present in BrM, G2, G3, and G4. **(B)** Subclone evolution Sc7->Ln9. Normalized AF of short variants on chromosome 14 clustered in C7 in red corresponding to new variants appeared in Sc7 shared by only Ln1 and Ln9. **(C)** Subclone evolution Sc5->Sc6->Sc9. The top panel is normalized AF of short variants on chromosome 14 clustered in C5, C6, and C9 depicted in red, green, and blue respectively. The bottom panel showed that BrP contains Sc5 which also appeared in BrM. BrM also contains Sc5's descendant subclone Sc6 which seeded Ln7. Sc9 existed in Ln1 and was the common ancestral clone seeding G2, G3 and G4. **(D)** Estimated CP based on normalized AF of each subclone in BrP, BrM, and Ln1.

Supplementary Figure S6

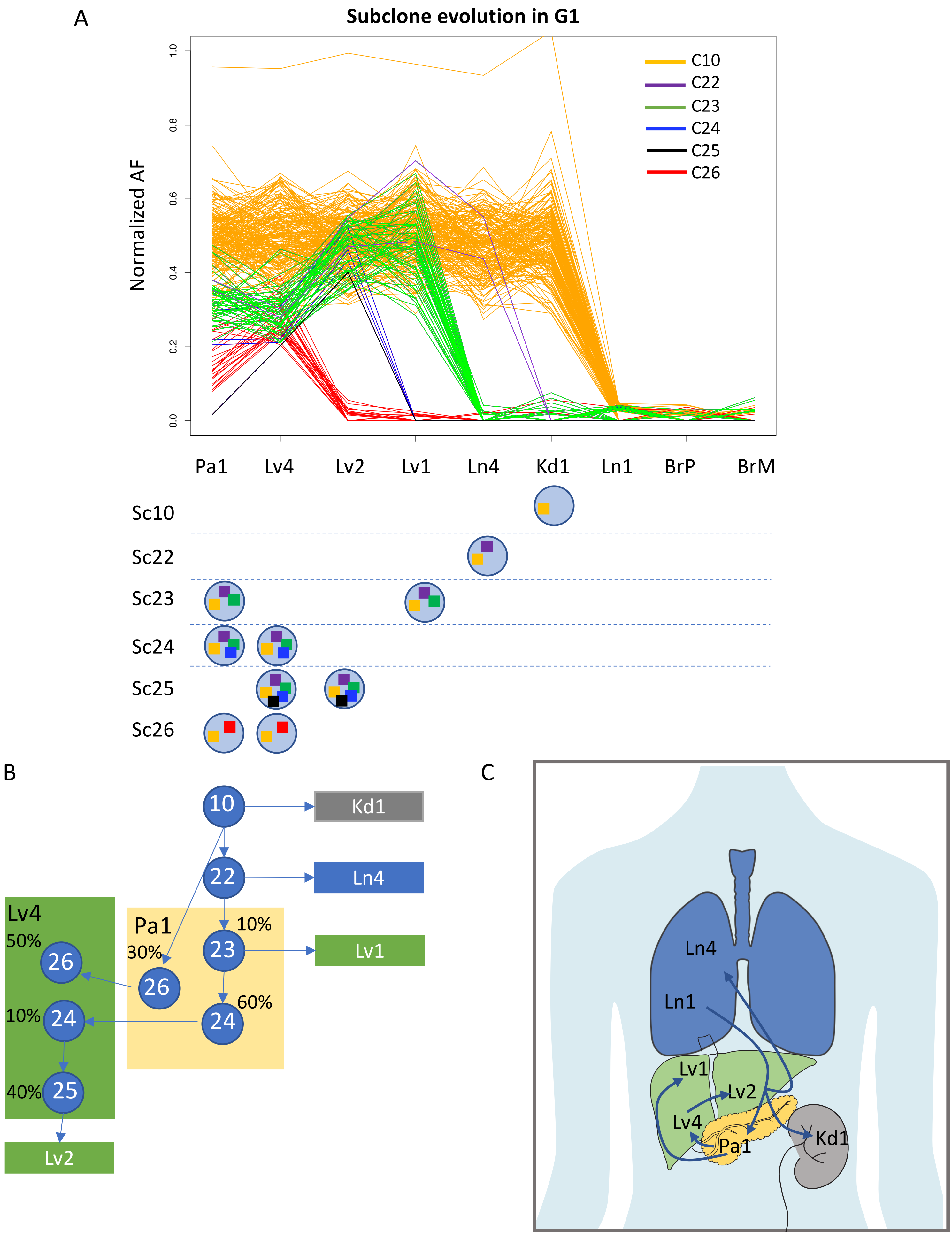

**Supplementary Figure S6. Subclone evolution and expansion for samples in G1. (A)** The top panel is normalized AF of G1 specific somatic short variants on chromosome 2, 7, 9, 11, 14, 15, 16, 21, and 22 clustered into C10, C22, C23, C24, C25, and C26 depicted in orange, purple, green, blue, black, and red respectively. The bottom panel shows which subclones each sample in G1 contains. Variants in C10 shared by all samples in G1 but not in Ln1, BrP and BrM. Sample Kd1, Ln4, Lv1 and Lv2 are clonal, whereas Lv4 and Pa1 had different subclones. Lv4 were polyclonal seeded by Pa1. **(B)** Estimated CP of each subclone of Pa1 and Lv4. Each blue circle represents an inferred subclone and each box represents a sample. **(C)** Migration pattern illustration for samples in G1.

Supplementary Figure S7

A

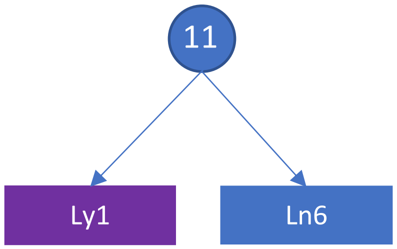


B

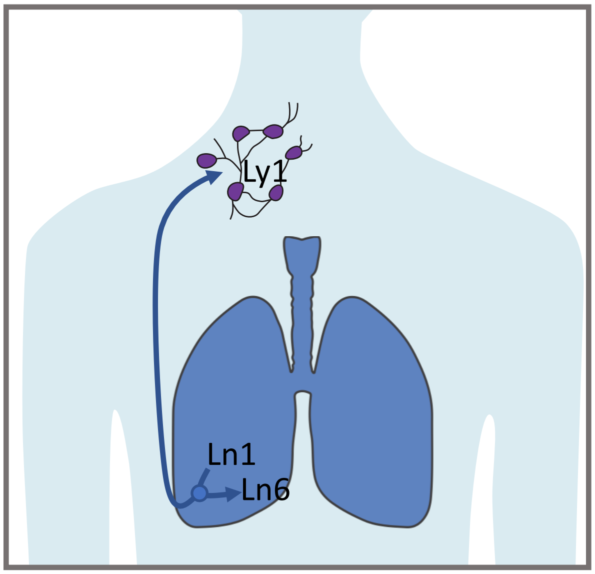

**Supplementary Figure S7. Samples in G2, Ly1 and Ln6 were clonal. (A)** G2 samples Ly1 and Ln6 were seeded by common ancestor Sc11. Each blue circle represents an inferred subclone and each box represents a sample. **(B)** Migration pattern illustration for samples in G2.

Supplementary Figure S8

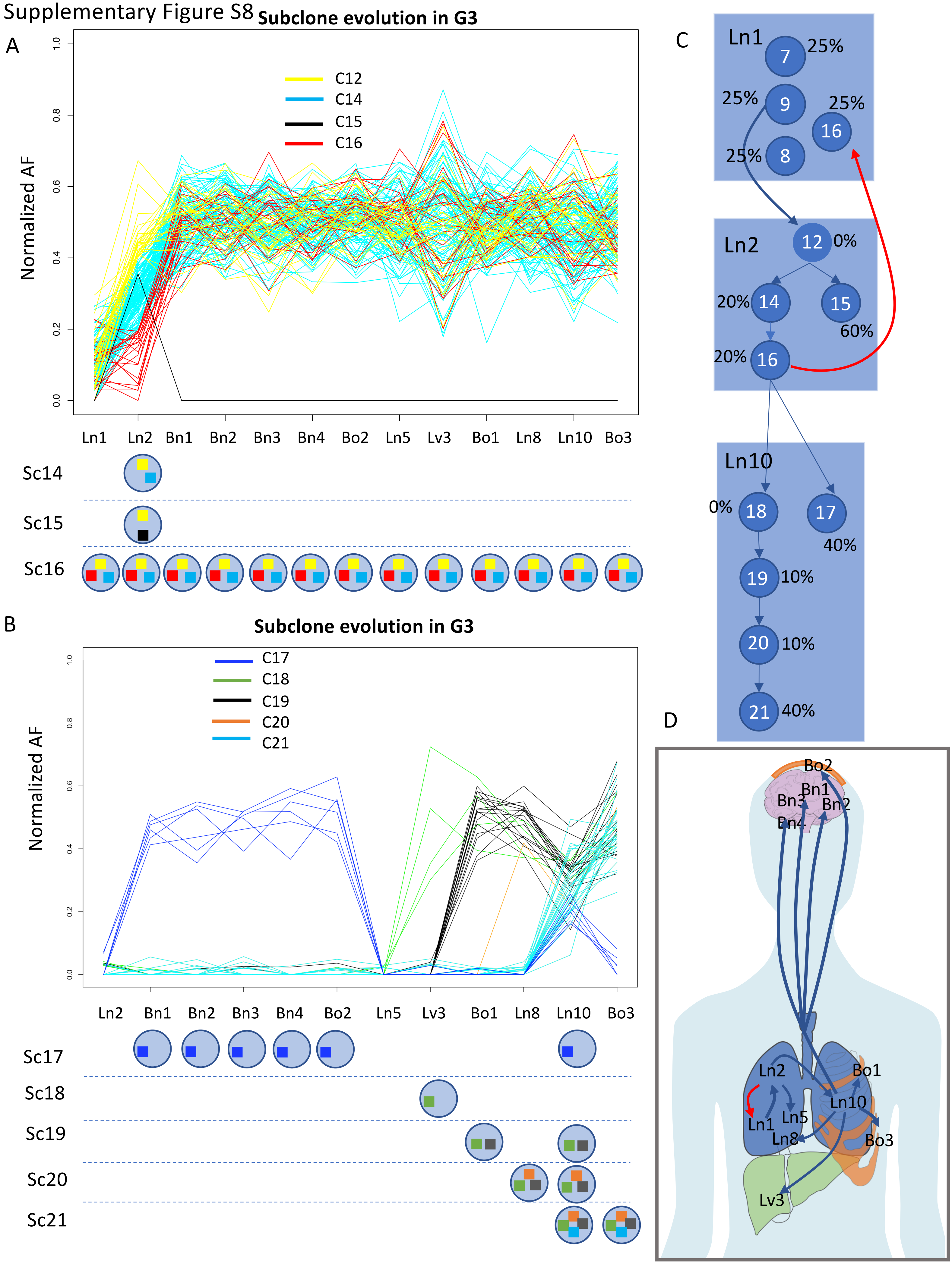

**Supplementary Figure S8. Subclone evolution and expansion for samples in G3. Normalized AF of G3 specific somatic short variants on chromosomes 2, 7, 9, 11, 14, 15, 16, 21, and 22. (A)** The top panel is normalized AF of short variants clustered into C12, C14, C15, and C16 depicted in yellow, cyan, black, and red respectively. The bottom panel is a schematic plot of which sample contains which subclones. The gradient AF of C12, C14, and C16 in Ln2 suggested evolution process Sc12->Sc14->Sc16 in Ln2. Thus the Ln2 was the birth-place of Sc16. The evidence that Sc16 also presented in Ln1 means that Sc16 recolonized Ln1 (red arrow in C). **(B)** The top panel is normalized AF of short variants clustered into C17, C18, C19, C20, and C21 depicted in blue, green, black, orange, and cyan respectively. The bottom panel is a schematic plot of which sample contains which subclones. Four brain samples and Bo2 evolved from a common ancestor Sc17 which also presented in Ln10. Sc18-Sc21 each of which seeded Lv3, Bo1, Ln8 and Bo3 respectively evolved in Ln10 indicating that Ln10 served as an incubator. **(C)** Estimated CP of each subclone of Ln2 and Ln10. Each blue circle represents an inferred subclone and each box represents a sample. **(D)** Migration pattern illustration for samples in G3.

Supplementary Figure S9

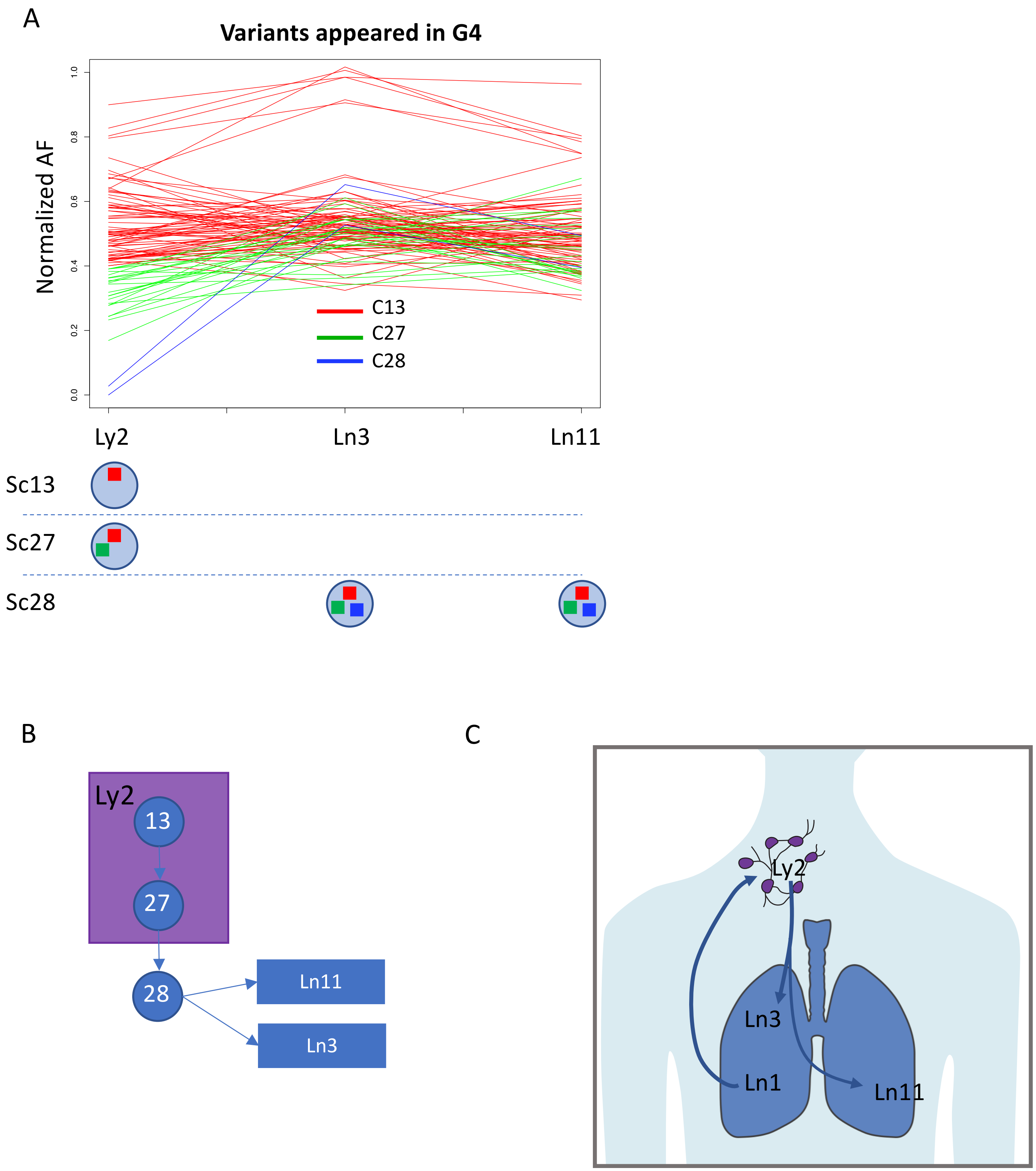

**Supplementary Figure S9. Subclone evolution and expansion for samples in G4. (A)** The top panel is normalized AF of G4 specific somatic short variants on chromosome 2, 7, 9, 11, 14, 15, 16, 21, and 22 clustered into C12, C14, C15, and C16 depicted in yellow, cyan, black, and red respectively. The bottom panel is a schematic plot of which sample contains which subclones. **(B)** Reconstructed subclone structure for Ly2 reveals that Ln11 and Ln3 were seeded by a further evolved subclone Sc28 from Ly2. Each blue circle represents an inferred subclone and each box represents a sample. **(C)** Migration pattern illustration for samples in G4.

#### Supplementary Figure S10

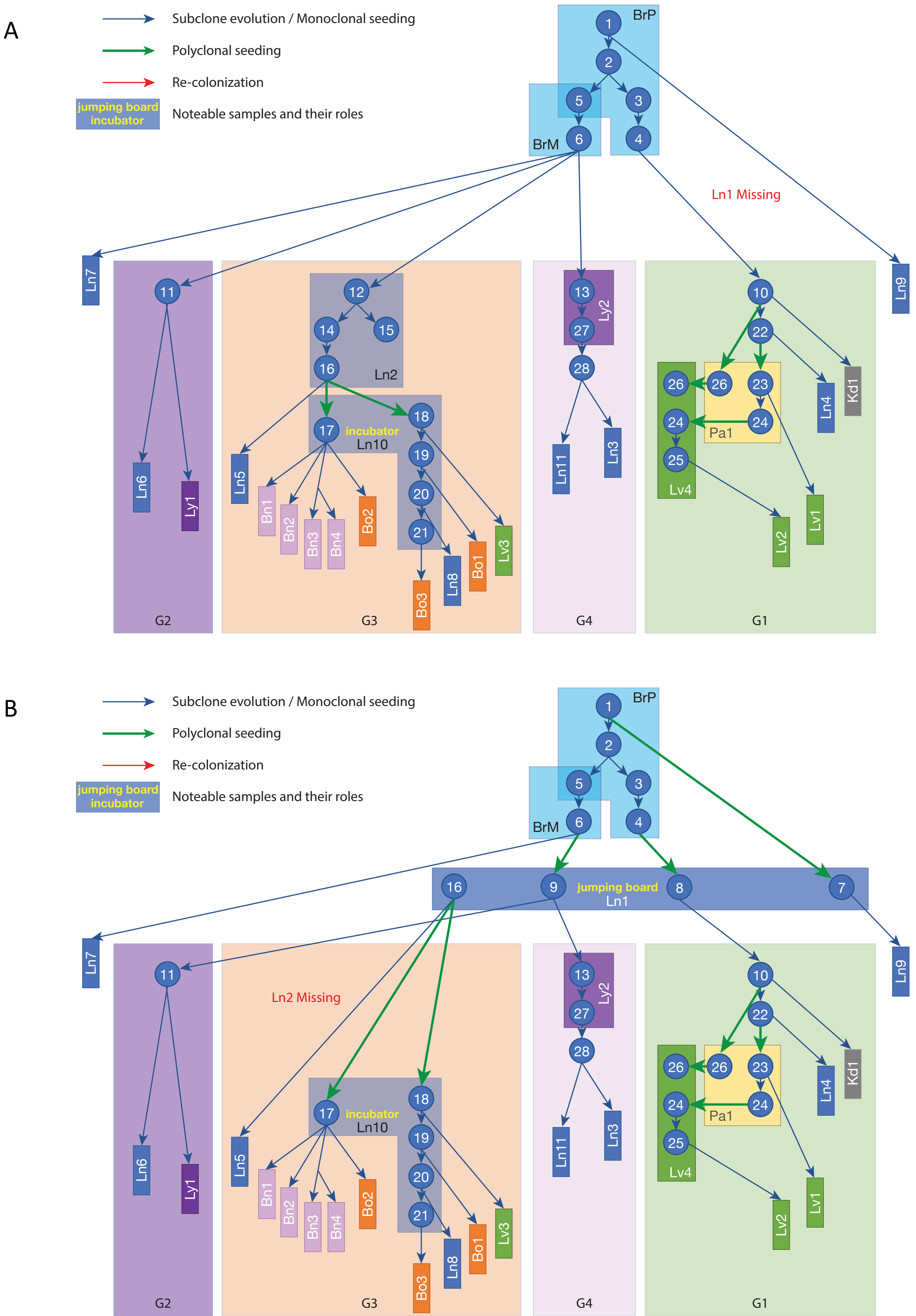

Supplementary Figure S10

C

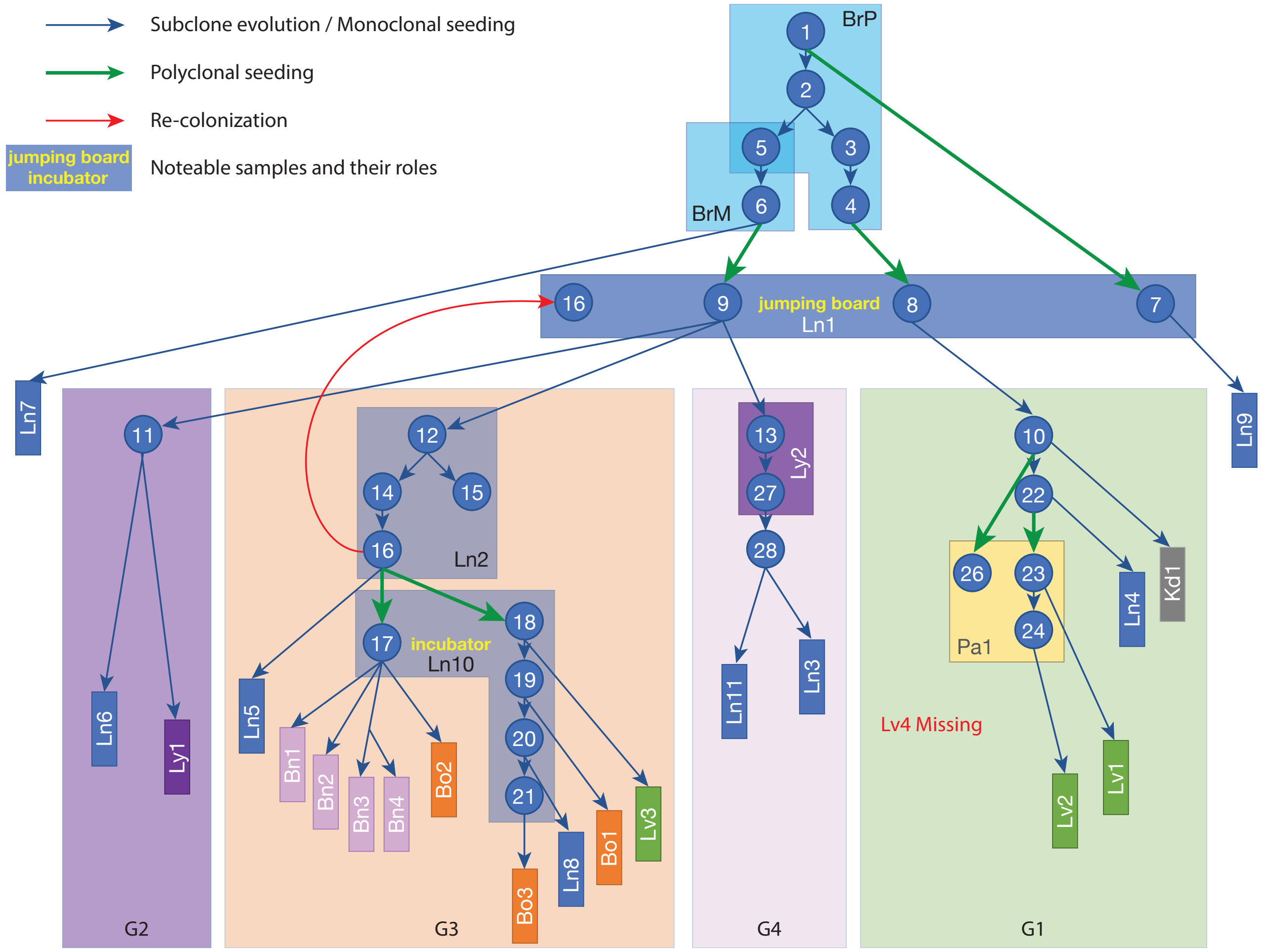

**Supplementary Figure S10. Alternative subclonal expansion scenario with missing samples Ln1 (A), Ln2 (B), Lv4 (C).**

Supplementary Figure S11

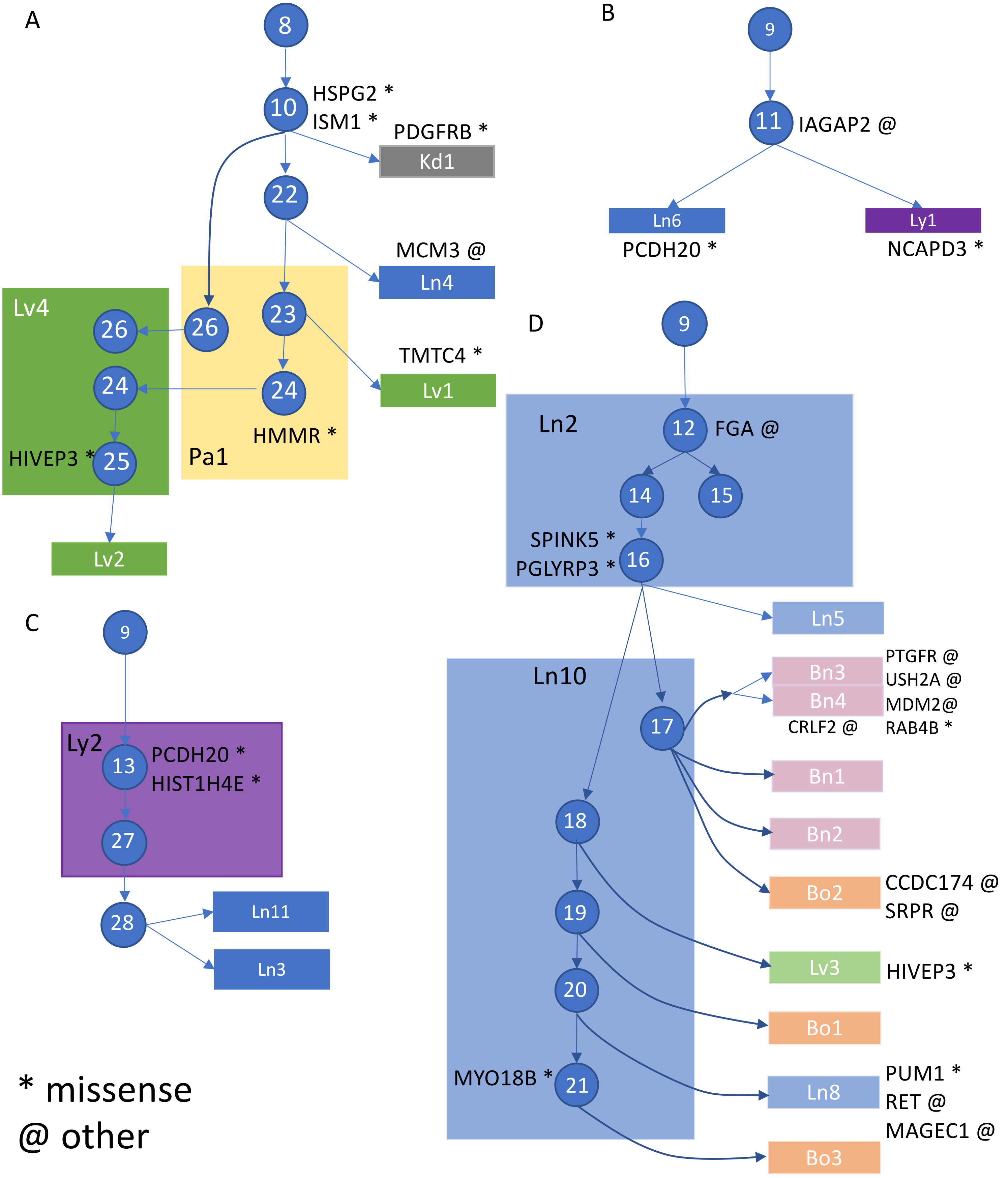


* missense
@ other

**Supplementary Figure S11. Group or sample specific somatic coding variants (A) G1, (B) G2, (C) G4, and (D) G3.** Missense mutations are depicted as *, and other coding mutations such as frameshift are depicted as @.

Supplementary Figure S12

A

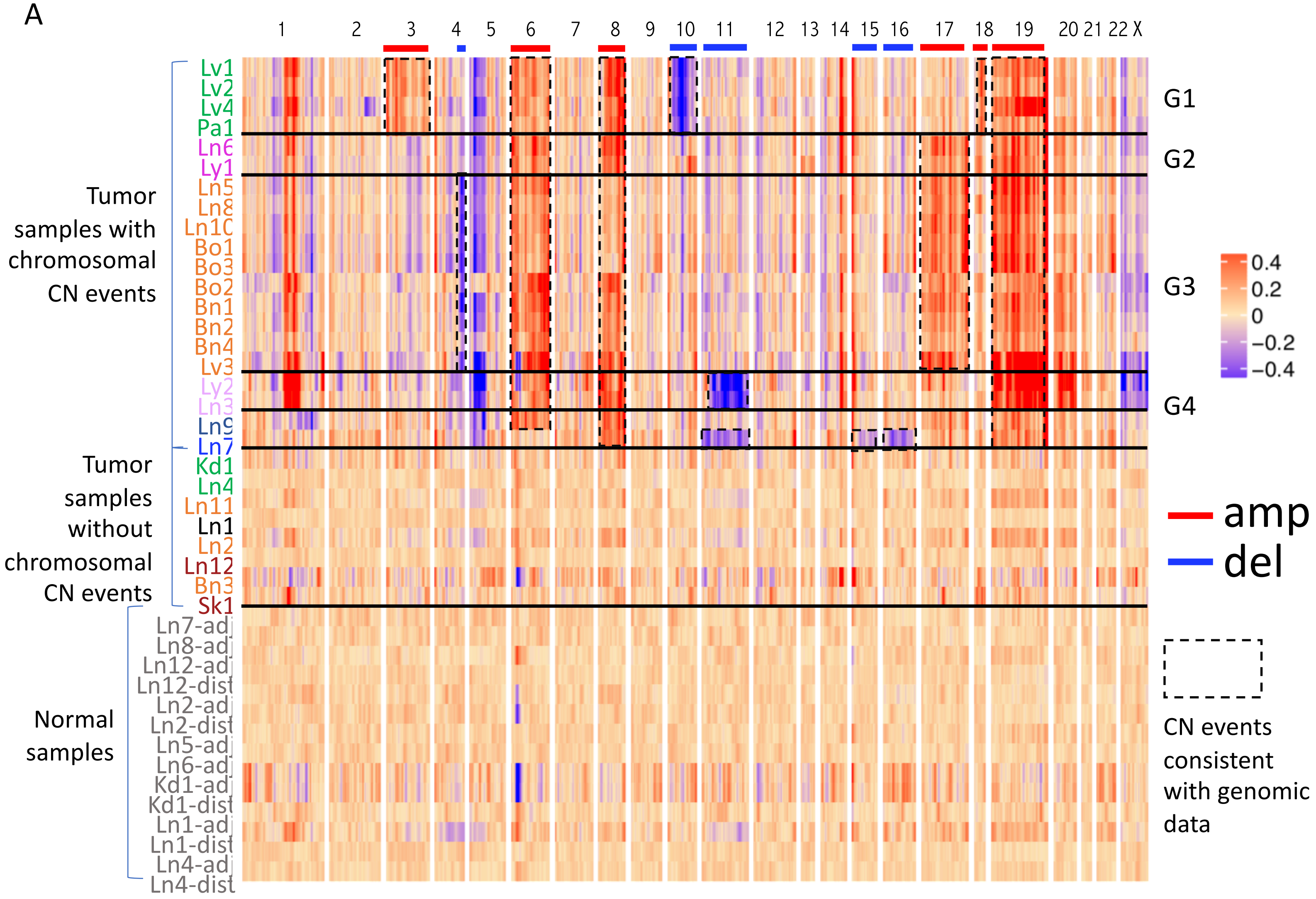


B

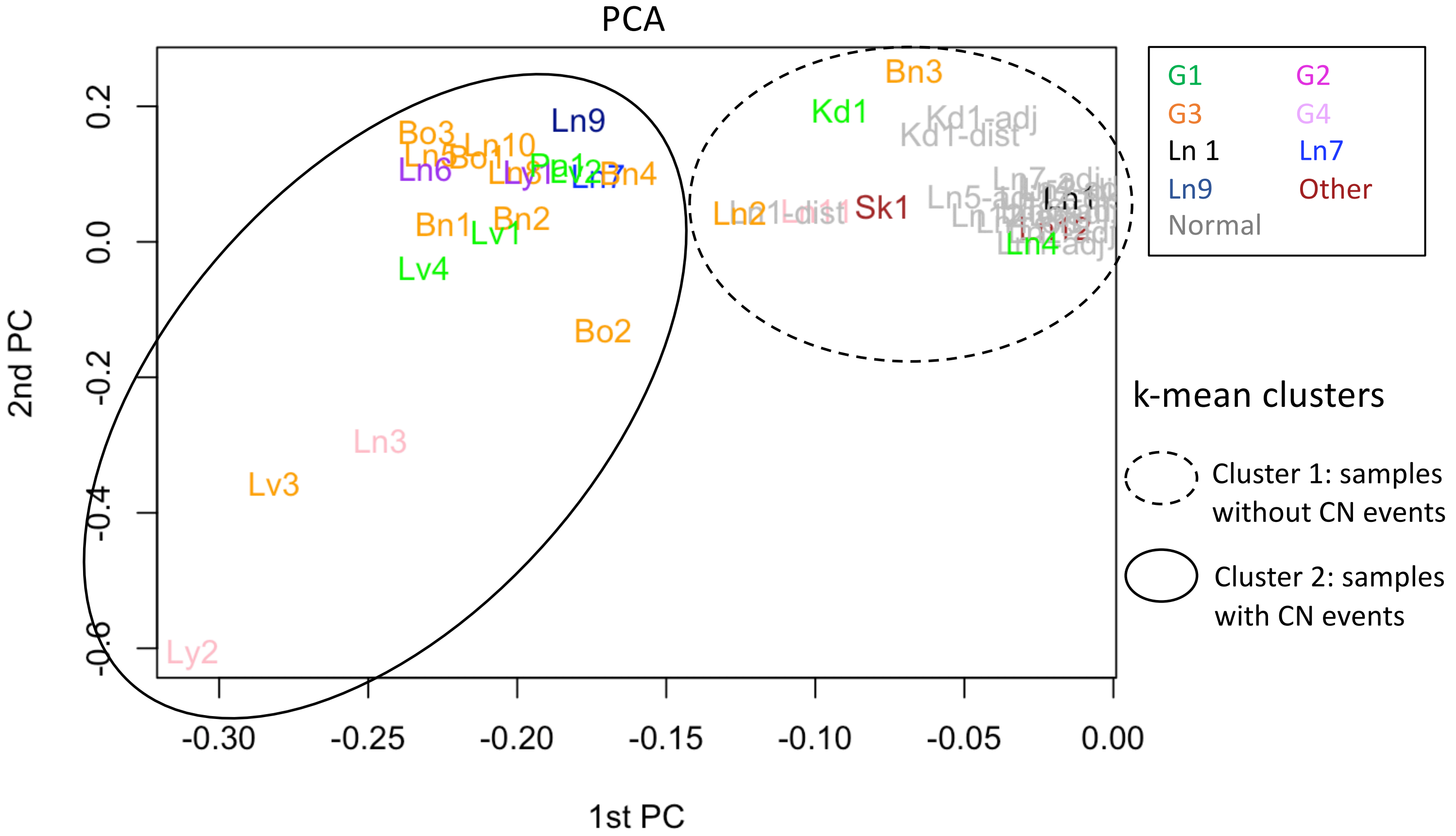

Supplementary Figure  S12

C

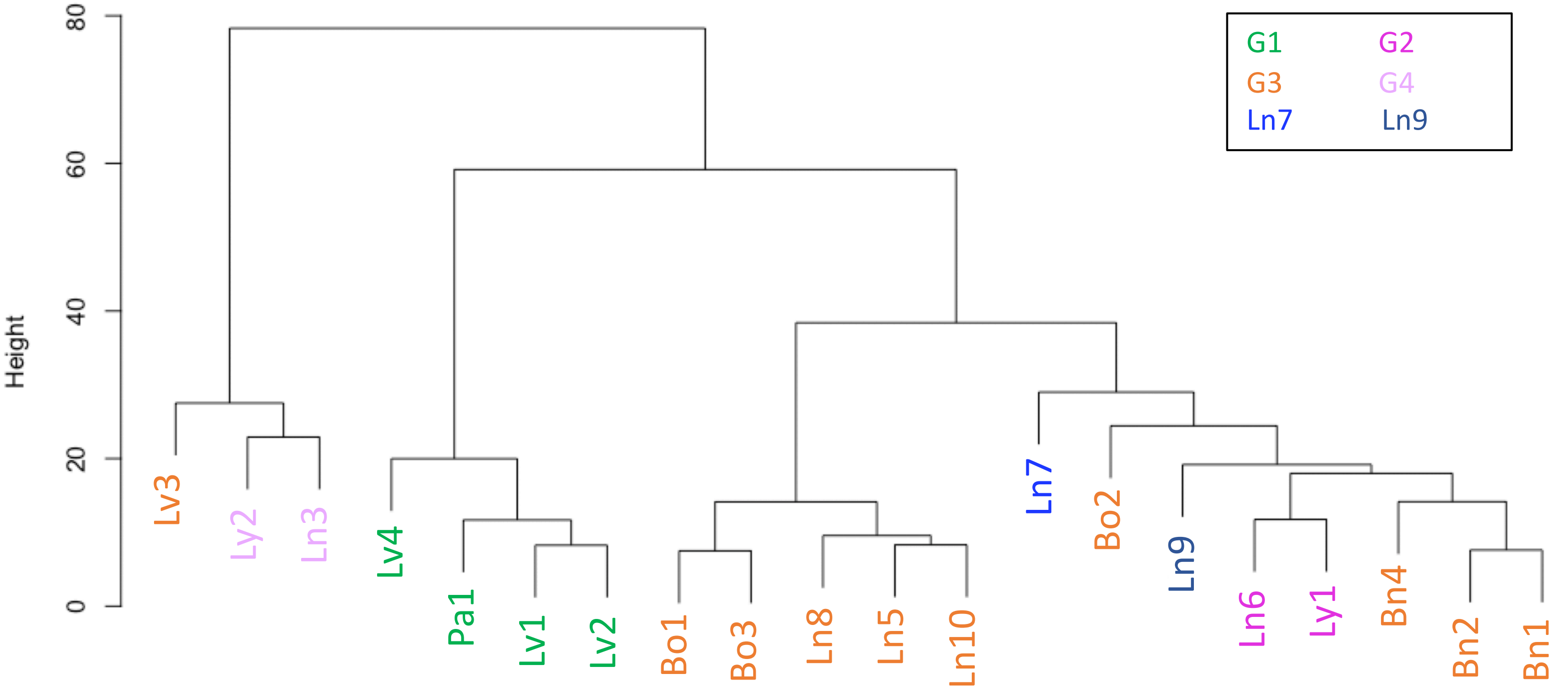

**Supplementary Figure S12. Copy number inferred from RNA sequencing data recapitulates genomic CNV events. (A)** Copy number profiles inferred from the expression fold-change averaged across 101-gene windows for all samples. Twenty tumor samples from the top showed significant CN changes and recapitulated the presence of large CNVs from the WGS data. Major CN events from WGS data are highlighted in dotted box. Red and blue lines indicate amplification and deletion events respectively. **(B)** PCA analysis and unsupervised k-means clustering on the expression fold-change averaged across 101-gene windows showed that 8 tumor samples (Ln1, Ln2, Ln11, Ln4, Kd1 and Bn3) clustered together with normal samples indicating that they had heavy normal tissue contamination. **(C)** Hierarchical clustering of the twenty purest tumor samples showed that most samples in the same group defined by genomic data also clustered together.

Supplementary Figure S13

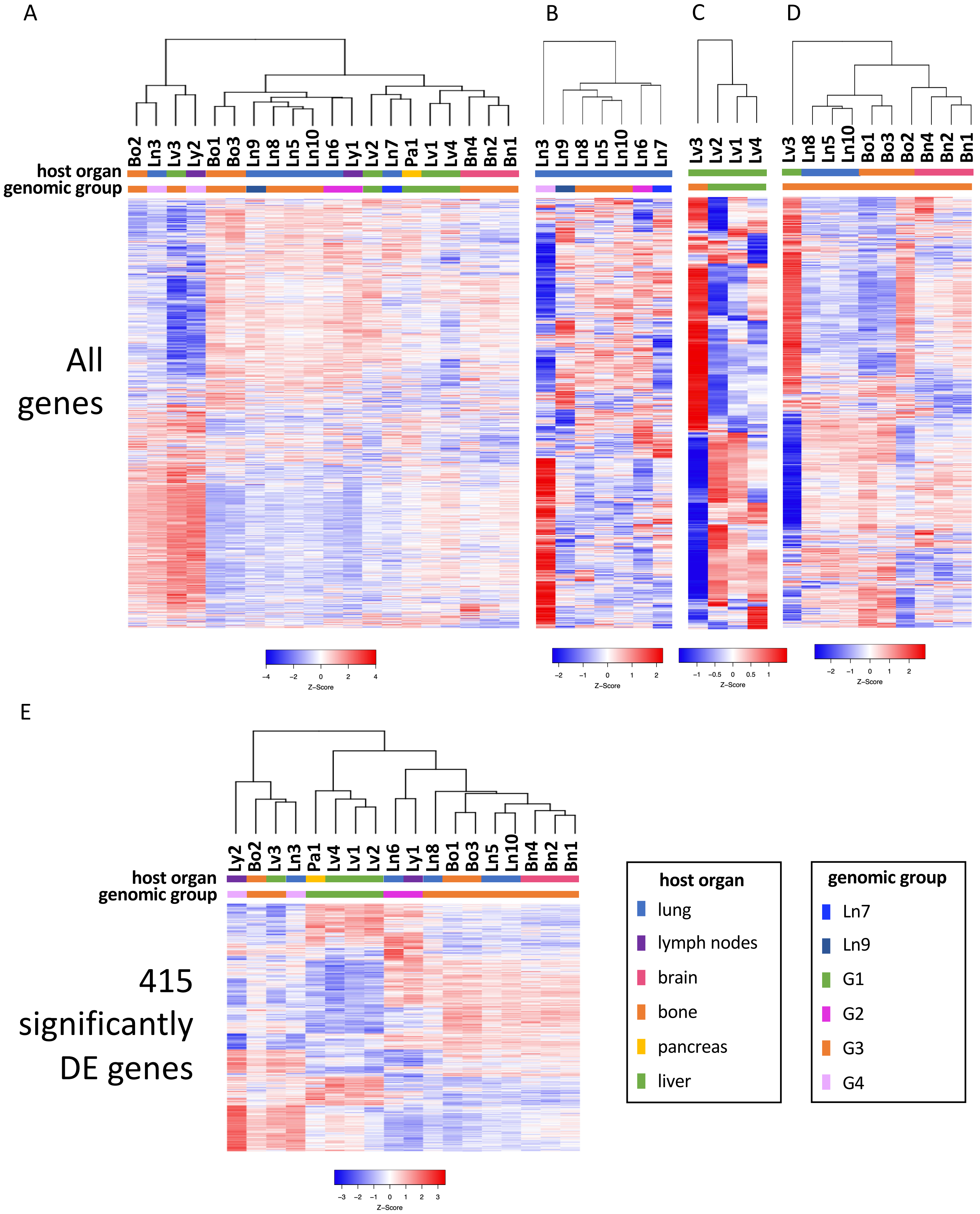

**Supplementary Figure S13. Transcriptomic profiling of metastatic tumors.** (A) Heatmap of gene expressions across 20 metastatic samples of high tumor purity. (B) Heatmap of gene expressions of 7 lung metastatic samples. Unsupervised clustering showed that Ln5, Ln8 and Ln10, which all belong to G3, were more similar to each other than with Ln6 of G2 and Ln3 of G4. (C) Heatmap of gene expressions of 4 liver metastatic samples. Unsupervised clustering showed that Lv1, Lv2 and Lv4, which all belong to G1, were more similar to each other than to Lv3 in G3. (D) Heatmap of gene expressions of 10 samples in G3 which contained the most diverse host organ types. Unsupervised clustering showed that samples located in the same organ tend to cluster together. (E) Heatmap of 415 significantly differentially expressed genes across samples in G1, G2, G3 and G4.

Supplementary Figure S14

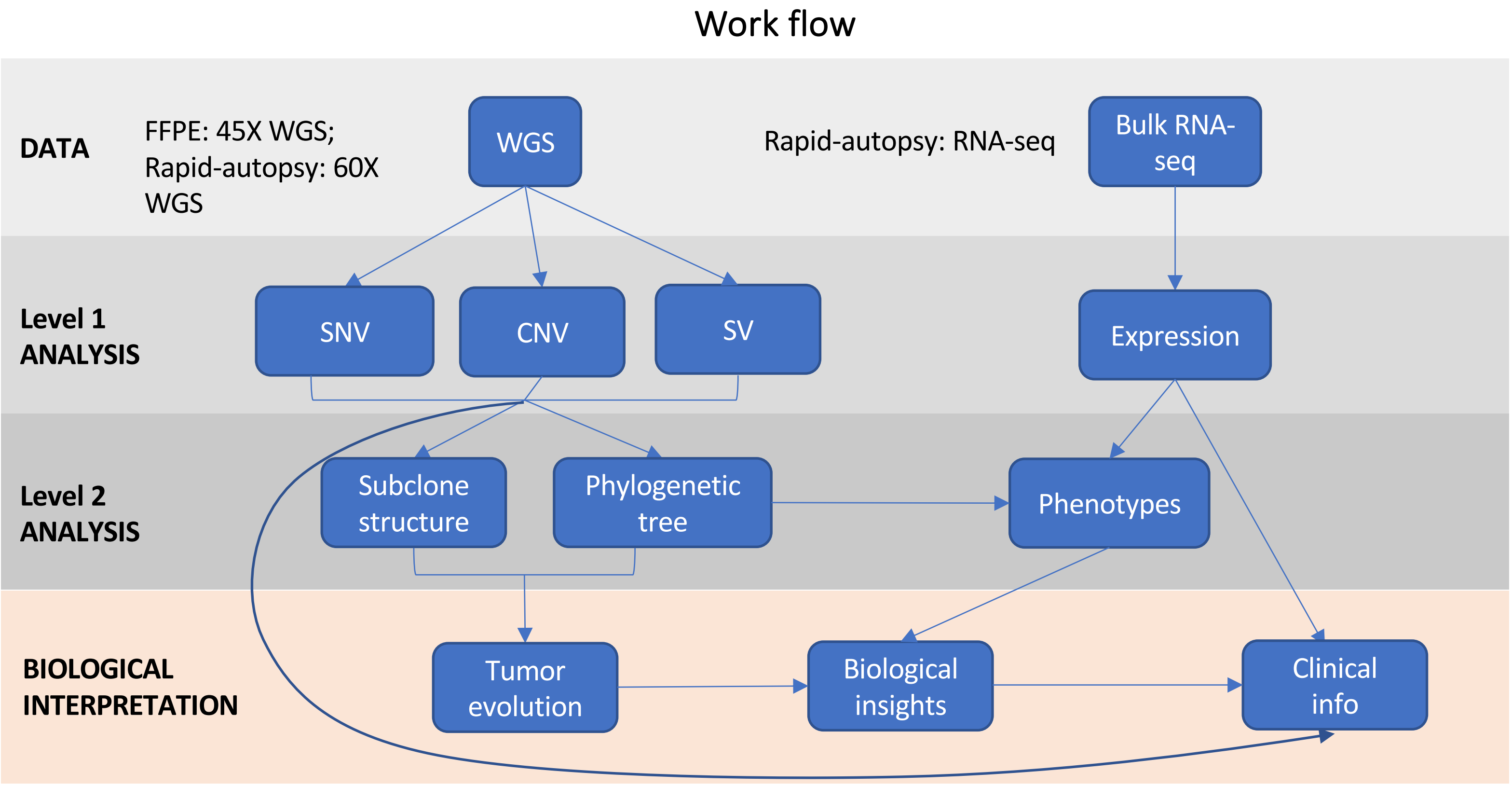

**Supplementary Figure S14. Overview of the workflow for this study.**

Supplementary Figure S15

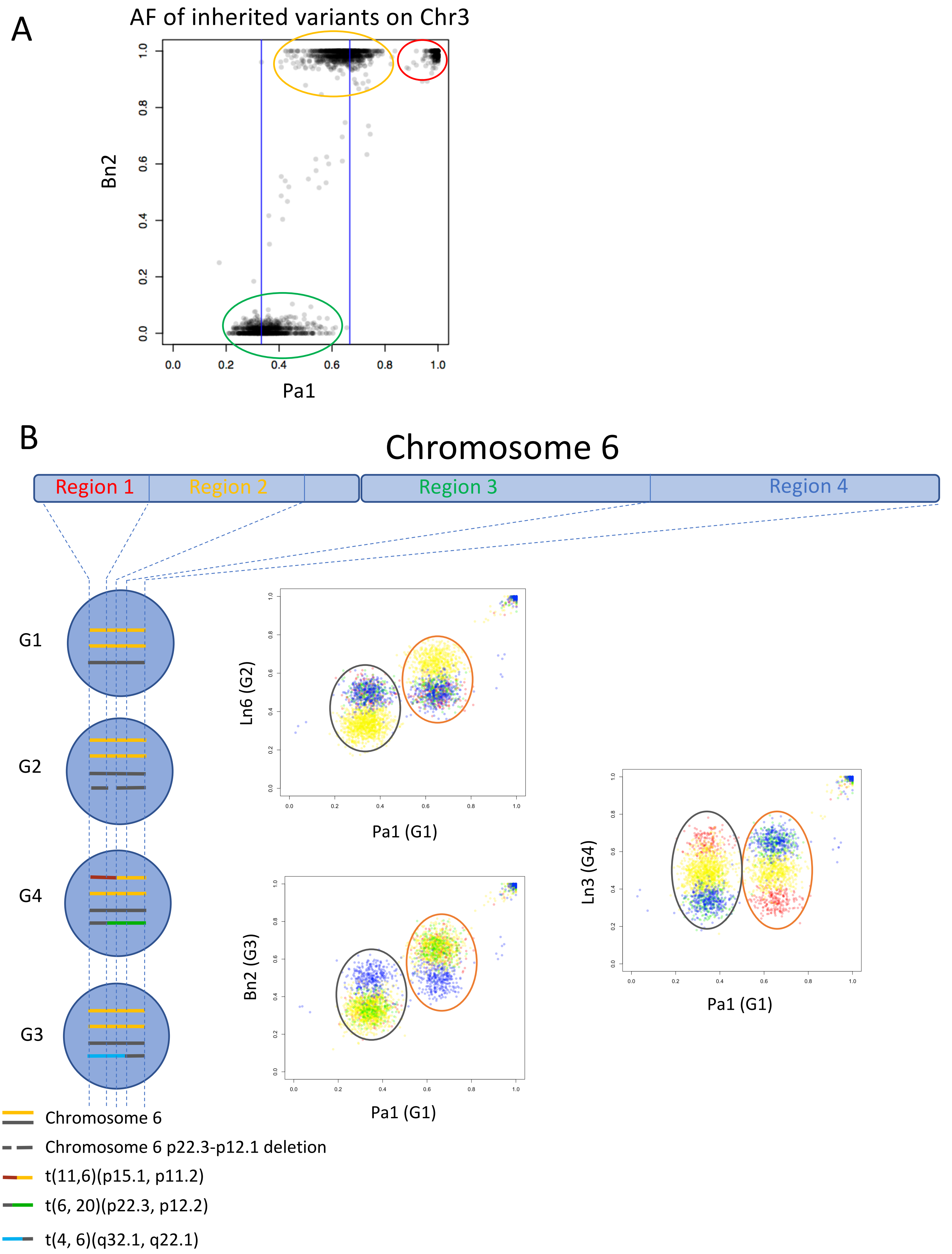

**Supplementary Figure S15. Allele specific CNV/LOH call. (A)** Scatter plot of allele frequencies of inherited variants on chromosome 3 between sample Pa1 and Bn2. **(B)** Allele frequencies of inherited variants on chromosome 6. Chromosome 6 was divided into four regions based on the breakpoints that caused different structural variants in our samples in different groups. Three scatter plots showed the allele frequencies of inherited variants in different regions between two samples. Each dot represents a variant. Color of the dots matches the region of the variant.

Supplementary Figure S16

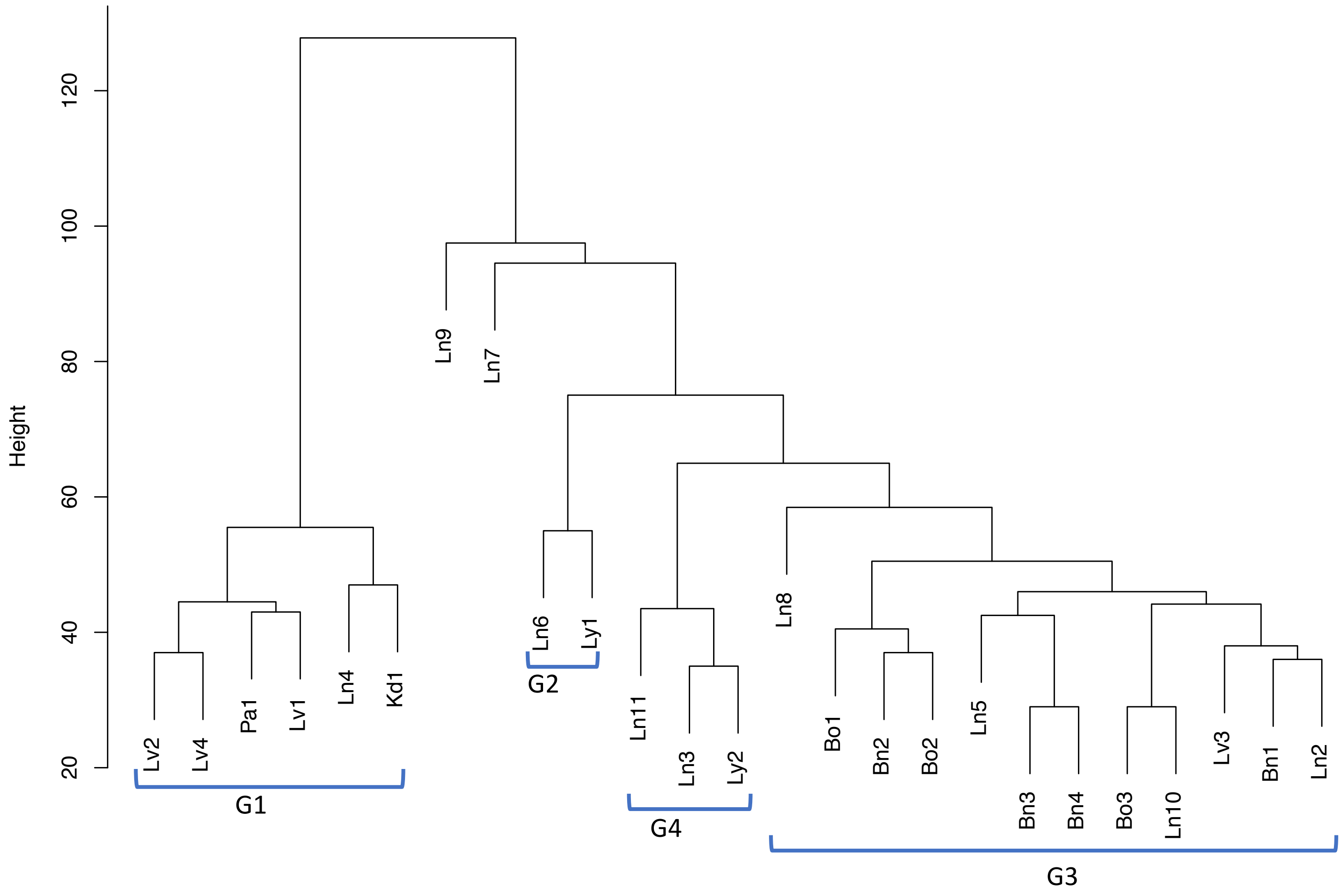

**Supplementary Figure S16. Automated sample clustering based on hamming distance between samples which were encoded as a vector of 0 (absence) or 1 (presence) of somatic short variants on chromosome 14.**

Supplementary Figure S17

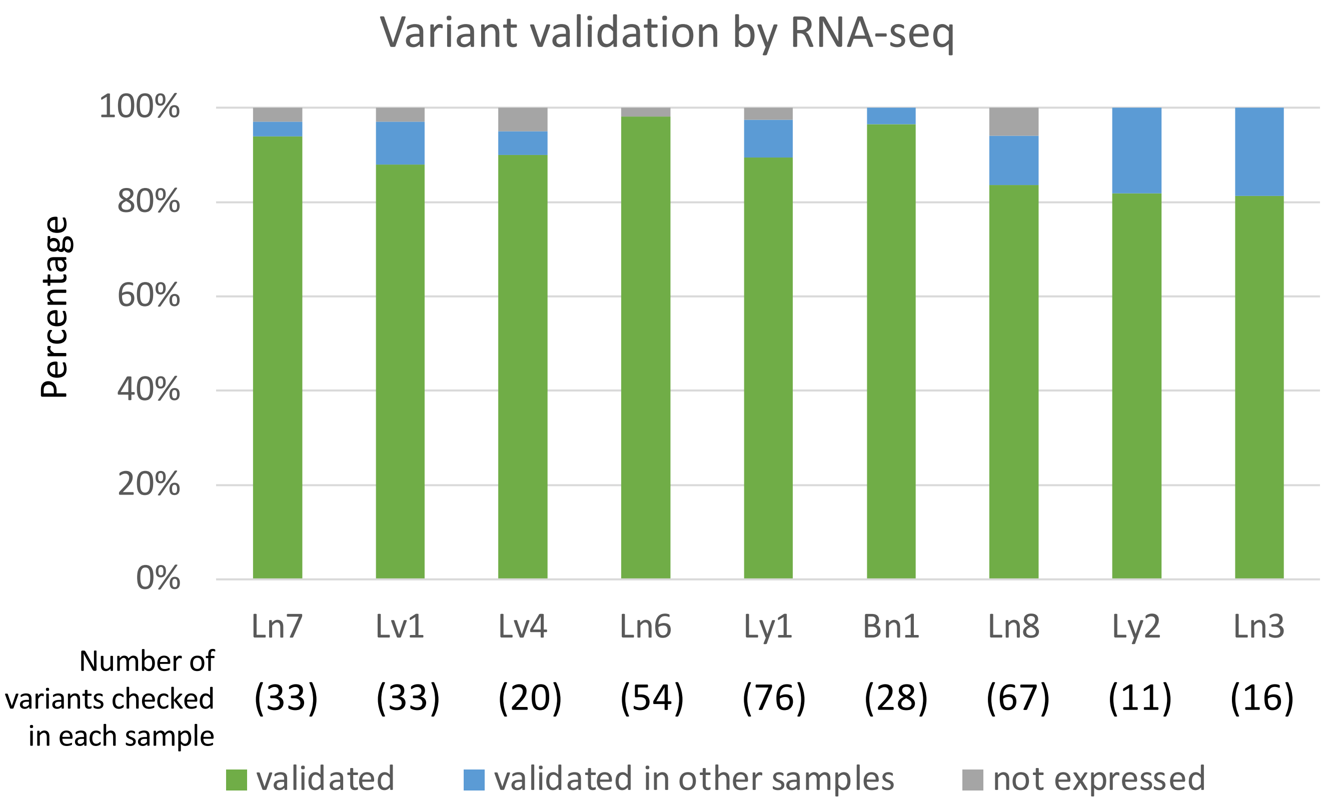

**Supplementary Figure S17. Somatic genomic short variants validation by RNA-seq data.** Two samples randomly picked from each group as well as Ln7 were used for somatic genomic variant validation. For any given tumor sample, only somatic short variants that have greater than 10% genomic VAF, and have at least 10 reads in the paired RNA-seq were considered. The number of variants checked in each sample is shown under the sample name. Green portion represents the percentage of variants that were validated in paired RNA-seq data in each sample. Blue portion represents the percentage of the variants that were validated in RNA-seq data of other samples which share the same variants. Grey portion represents the variants that were not detected in either paired RNA-seq data or other samples' RNA-seq data. More than 90% of variants can be either "validated" or "validated in other samples".
